# Tvp38/DedA Family Protein mediates biogenesis of chloroplast peripheral reticulum and envelope-to-thylakoid contacts in *Arabidopsis*

**DOI:** 10.64898/2026.10.13.758368

**Authors:** Melanie Voelkel-Lambertz, Michał Bykowski, Anja Roediger, Khushbu Kumari, Christopher Grefen, Ralph Bock, Sacha Baginsky, Łucja Kowalewska, Omar Sandoval-Ibáñez

## Abstract

Chloroplasts of land plants are dynamic organelles essential for photosynthesis, metabolism, and plant adaptation to changing environmental conditions. Within chloroplasts, the inner envelope and thylakoid membranes are spatially separated but functionally interconnected. The inner envelope harbors enzymes involved in lipid synthesis for internal membranes and controls the exchange of metabolites and proteins, whereas the thylakoid membranes accommodate the machinery of the photosynthetic electron transport chain responsible for converting light energy into chemical energy. Therefore, efficient communication between these membrane systems is essential for maintaining chloroplast homeostasis and sustaining photosynthetic activity.

In this study we have characterized the role of Tvp38/DedA Family Protein (TVPFP) in mediating inner envelope-to-thylakoid communication in *Arabidopsis thaliana*. Despite exhibiting wild-type-like growth and similar accumulation of the major thylakoid-anchored photosynthetic complexes, *tvpfp* mutants displayed reduced photosynthetic activity and increased abundance of photorespiration- and pigment-related proteins. Our results revealed that TVPFP localizes to the chloroplast envelope and undergoes post-translational modification when localized to the thylakoids. The unmodified form of TVPFP is enriched in high-molecular-weight complexes, suggesting that TVPFP oligomers are partially dissolved upon thylakoid relocaltion. Transmission electron microscopy data showed reduced abundance of vesicles as well as chloroplast peripheral reticulum (CPR) structures connecting the envelope and thylakoids in *tvpfp* mutants. Together, our results demonstrate that TVPFP fine-tunes photosynthesis by regulating vesicle abundance and biogenesis of CPR structures. Our findings shed light on the importance of envelope-to-thylakoid communication and the role of CPR structures in land plants.

## Introduction

The envelope of chloroplasts consists of an outer and an inner membrane that enclose a soluble compartment known as the stroma and an internal membrane system known as the thylakoids. While carbon fixation takes place in the stroma, the thylakoids house the photosynthetic machinery responsible for the light-dependent reactions, including photosystem II (PSII), the cytochrome *b*_6_*f* complex (Cyt*b*_6_*f*), photosystem I (PSI), and the chloroplast ATP synthase (cpATPase) (1). These complexes, embedded within the thylakoid membranes, are composed of numerous protein subunits, pigments, and cofactors that are essential for photosynthetic electron transfer (2, 3). Key steps in the assembly of the thylakoid membranes, including incorporation of the major structural lipids monogalactosyldiacylglycerol (MGDG) and digalactosyldiacylglycerol (DGDG) take place in the envelope membrane of the chloroplast (4). Therefore, macromolecular transport from the envelope to the thylakoid membranes is essential for chloroplast development (5).

Vesicles are one of the transport systems that contribute to envelope-to-thylakoid communication (6). While some proteins contributing to vesicle accumulation have been identified (7), only few mutants affected in vesicle formation exhibit abnormal phenotypes (6). Among them, the lipid-binding chloroplast-localized Sec14-like protein (CPSFL1) was identified as an essential factor required for vesicle accumulation (8). Genetic analyses of *cpsfl1* mutants and CPSFL1-overexpressing lines revealed that CPSFL1 abundance correlates with the accumulation of vesicles in chloroplasts (8).

Another mechanism facilitating communication between the envelope and the thylakoids involves direct membrane contact sites. Recent structural analyses have identified contact sites between the inner envelope membrane and the thylakoids in many photosynthetic organisms, including cyanobacteria, algae, and plants (9–11). However, the molecular components mediating the formation of contact sites between the envelope membrane and the thylakoid membranes remain largely unknown.

De-etiolation experiments provide a robust system for studying chloroplast development (12) and enable the discovery of new plastid-localized proteins that participate in thylakoid biogenesis and assembly of the photosynthetic machinery. By leveraging a de-etiolation system (13), here we have identified and characterized the molecular function of the plastid-localized Tvp38/DedA Family Protein (TVPFP) in *Arabidopsis thaliana*. Our findings reveal reduced photosynthetic activity at PSII and increased accumulation of both photorespiration- and pigment-related proteins in *tvpfp* mutants. TVPFP localizes to the chloroplast envelope fraction and is present in high molecular weight complexes. In addition, TVPFP undergoes post-translational modification when localized to the grana margins within the thylakoids, and is then found exclusively as monomer and dimer. The *tvpfp* mutants display low accumulation of structures known as chloroplast peripheral reticulum (CPR). Although the role of CPR is poorly understood, an inverse correlation between abundance of CPR and photorespiration has been reported (14, 15). Our results show that CPR structures connect the envelope with the thylakoid membranes. The reduced CPR contents in *tvpfp* mutants are in agreement with the high accumulation of photorespiration-related proteins. Overall, our study assigns a role to TVPFP in the biogenesis of CPR structures and sheds light on the importance of the communication between the envelope membrane and the thylakoid membranes.

## Results

### Identification of TVPFP as candidate for chloroplast development

To identify new factors contributing to chloroplast development, we screened a time-resolved RNA-seq dataset of de-etiolation in tobacco leaves (13). We identified the transcript encoded by locus 104241569 in *Nicotiana sylvestris,* one of the diploid progenitor species of the allotetraploid tobacco *Nicotiana tabacum,* as a potential candidate gene. Transcripts from this gene progressively increased during the early stages of greening, reaching a 5-fold increase relative to dark-grown seedlings after 120 min of illumination (Fig. S1). The putative ortholog in *Arabidopsis thaliana* corresponds to AT1G22850, which is highly expressed in cotyledons and young leaves (Fig. S2). AT1G22850 is annotated as <u>Tvp</u>38 <u>F</u>AMILY <u>P</u>ROTEIN (TVPFP) and belongs to the GreenCut2 gene set, which comprises genes conserved throughout the Viridiplantae lineage and is represented by orthologs in cyanobacteria, algae, and plants (16). The locus encodes a putative chloroplast-localized protein containing a predicted chloroplast transit peptide (cTP) at the N-terminus (17), five predicted transmembrane domains, and a VMP1/TMEM41/TVP38 (VTT) domain toward the C-terminus (Fig. S3A). Similar to locus 104241569 in tobacco, *TVPFP* transcript levels increased upon de-etiolation, as revealed by our RT-PCR analysis (Fig. S3B). To investigate the role of TVPFP in chloroplast development, we screened collections of T-DNA insertion lines for mutants in AT1G22850. We identified a line harboring a T-DNA insertion in the third exon (*tvpfp-1*; SALK_095443; Fig. 1A) and isolated homozygous mutants (Fig. S3C). An additional mutant was generated by CRISPR/Cas9-mediated genome editing. A pair of single-guide RNAs (sgRNAs) was designed to target the first exon, resulting in deletion of the entire chloroplast transit peptide (*tvpfp-2*; Fig. 1A). Genotyping and genomic DNA sequencing confirmed the homozygous state of *tvpfp-2* and the deletion of 446 bp, corresponding to the first 144 amino acids of the encoded protein (Fig. S3D). The absence of detectable TVPFP transcripts in *tvpfp-1* and the strongly reduced transcript levels in *tvpfp-2* (Fig. S3E) indicate that *tvpfp-1* is a knockout mutant, whereas *tvpfp-2* represents a strong knockdown allele lacking a functional chloroplast-targeting sequence.

**Figure 1.**
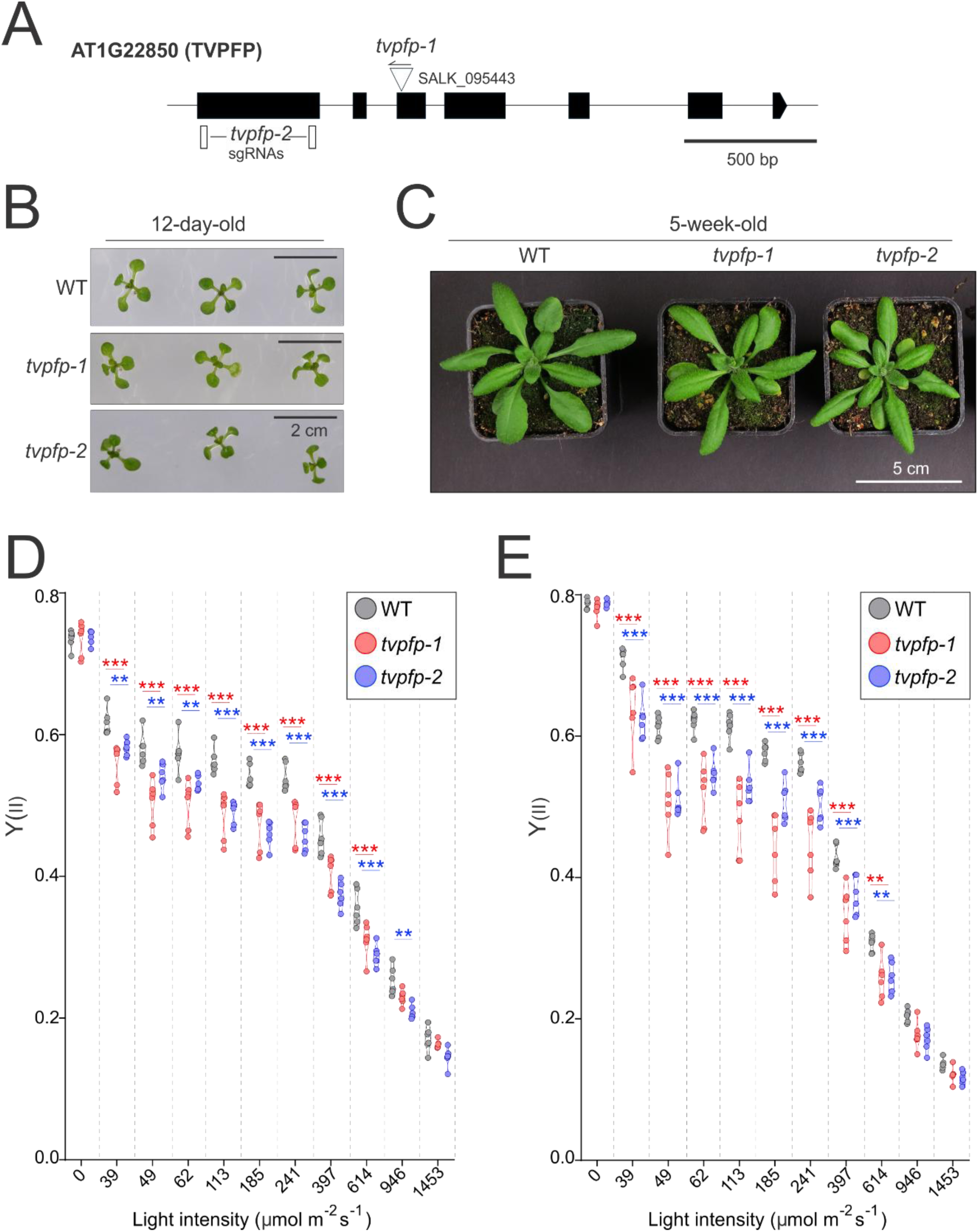
Phenotypic characterization of *tvpfp* mutants. **A)** Schematic representation of the *AT1G22850* (*TVPFP*) locus. The figure shows the position of the T-DNA insertion in *tvpfp-1* and the single-guide RNAs targeting the first exon in the *tvpfp-2* mutant. **B)** Phenotype of 12-day-old wild type (WT), *tvpfp-1*, and *tvpfp-2* plants grown under mixotrophic conditions. Scale bar = 2 cm. **C)** Phenotype of 5-week-old wild type (WT), *tvpfp-1*, and *tvpfp-2* plants grown under photoautotrophic conditions. Scale bar = 5 cm. **D–E)** Light-response curves of the quantum yield of photosystem II (Y_(II)_) in **D)** 12-day-old wild type (WT), *tvpfp-1*, and *tvpfp-2* plants grown under mixotrophic conditions and **E)** 5-week-old wild type (WT), *tvpfp-1*, and *tvpfp-2* plants grown under photoautotrophic conditions. Statistical differences between the mutants and WT were assessed using two-way ANOVA followed by Dunnett’s post hoc test. * *P* < 0.05, ** *P* < 0.01, *** *P* < 0.001. n = 6 biological replicates.

### TVPFP is not essential for plant development but fine-tunes photosynthesis

The increased accumulation of TVPFP transcripts during de-etiolation prompted us to analyze the phenotypes of the *tvpfp-1* and *tvpfp-2* mutants during greening (Fig. S3F-I). Similar to wild-type seedlings, the *tvpfp* mutants exhibited the characteristic apical hook and closed cotyledons of etiolated seedlings, suggesting that skotomorphogenesis is not affected by mutations in TVPFP (Fig. S3F). Upon illumination, wild type, *tvpfp-1*, and *tvpfp-2* seedlings underwent greening with similar kinetics, and no noticeable differences were observed between the genotypes after 6, 12, or 24 h of de-etiolation (Fig. 3G-I). Despite the light-induced upregulation of TVPFP transcript levels, these results indicate that TVPFP is not essential for chloroplast biogenesis during de-etiolation.

We next analyzed the phenotypes of *tvpfp-1* and *tvpfp-2* plants grown under mixotrophic conditions for 12 days, or autotrophic conditions for 4 weeks (Figs. 1B, C). Under mixotrophic conditions, both mutants exhibited size and morphology comparable to those of wild-type plants (Fig. 1B). Likewise, no phenotypic differences were observed between *tvpfp-1*, *tvpfp-2*, and wild-type plants grown under autotrophic conditions (Fig. 1C).

The absence of obvious developmental phenotypes prompted us to analyze PSII photosynthetic performance by pulse-amplitude modulation (PAM) fluorometry in wild type, *tvpfp-1* and *tvpfp-2* plants grown under mixotrophic or autotrophic conditions (Fig. 1D, E). Light-response curves of plants grown under mixotrophic conditions revealed a lower effective quantum yield of PSII (which reflects the proportion of photochemically active PSII reaction centers; Y_(II)_) in *tvpfp* mutants over a range of light intensities from 39 to 614 µmol photons m^-2^ s^-1^. Similar results were obtained under autotrophic conditions, where *tvpfp-1* and *tvpfp-2* mutants also displayed reduced Y_(II)_ compared to wild-type plants across the same range of light intensities.

The *tvpfp* mutants exhibited higher Y(NO) (which reflects the fraction of light energy passively dissipated by PSII) under low light intensities in both growth conditions (39 and 49 µmol photons m^-2^ s^-1^ under mixotrophic and autotrophic conditions, respectively) (Fig. S4A, B). Under mixotrophic conditions, *tvpfp-1* and *tvpfp-2* also showed slightly elevated Y(NO) at high light intensities (946 and 1453 µmol photons m^-2^ s^-1^), whereas under autotrophic conditions, Y(NO) values were comparable to those of the wild type at higher irradiances.

Consistent with these observations, *tvpfp* mutants exhibited higher non-photochemical quenching (NPQ), reflecting increased dissipation of excess excitation energy as heat, than wild-type plants under low light intensities (49 and 39 µmol photons m^-2^ s^-1^ under mixotrophic and autotrophic conditions, respectively) (Fig. S4C, D). Compared to the wild type, elevated NPQ values persisted in *tvpfp-1* and *tvpfp-2* up to 397 µmol photons m^-2^ s^-1^ under mixotrophic conditions and up to 614 µmol photons m^-2^ s^-1^ under autotrophic conditions.

The *tvpfp-1* and *tvpfp-2* mutants exhibited lower values for qL (which reflects the fraction of open reaction centers occupied at the Qa site) at low light intensity (39 µmol photons m^-2^ s^-1^) in both growth conditions (Fig. S4E, F). Collectively, these results suggest that TVPFP is required for fine-tuning of photosynthesis.

Since autotrophic cultivation may increase photosynthetic pressure and lead to indirect pleiotropic effects, we used 12-day-old plants grown under mixotrophic conditions for the subsequent characterization of TVPFP.

### *tvpfp* mutants overaccumulate proteins related to pigment synthesis and photorespiration

We analyzed the accumulation of diagnostic proteins from the main photosynthetic complexes in total protein extracts from wild type, *tvpfp-1*, and *tvpfp-2* plants grown under mixotrophic conditions by polyacrylamide (PAA) gel electrophoresis followed by immunodetection (Fig. 2A). Both *tvpfp* mutants exhibited wild type-like accumulation patterns of PSI (PsaA and PSAD) and its associated antenna complex LHCI (LHCA1), PSII (PsbD and PsbC) and its associated antenna complex LHCII (LHCB2), the cytochrome *b*_6_*f* complex (PetA), and the chloroplast-localized ATP synthase complex (AtpB and ATPC) (Fig. 2A).

**Figure 2.**
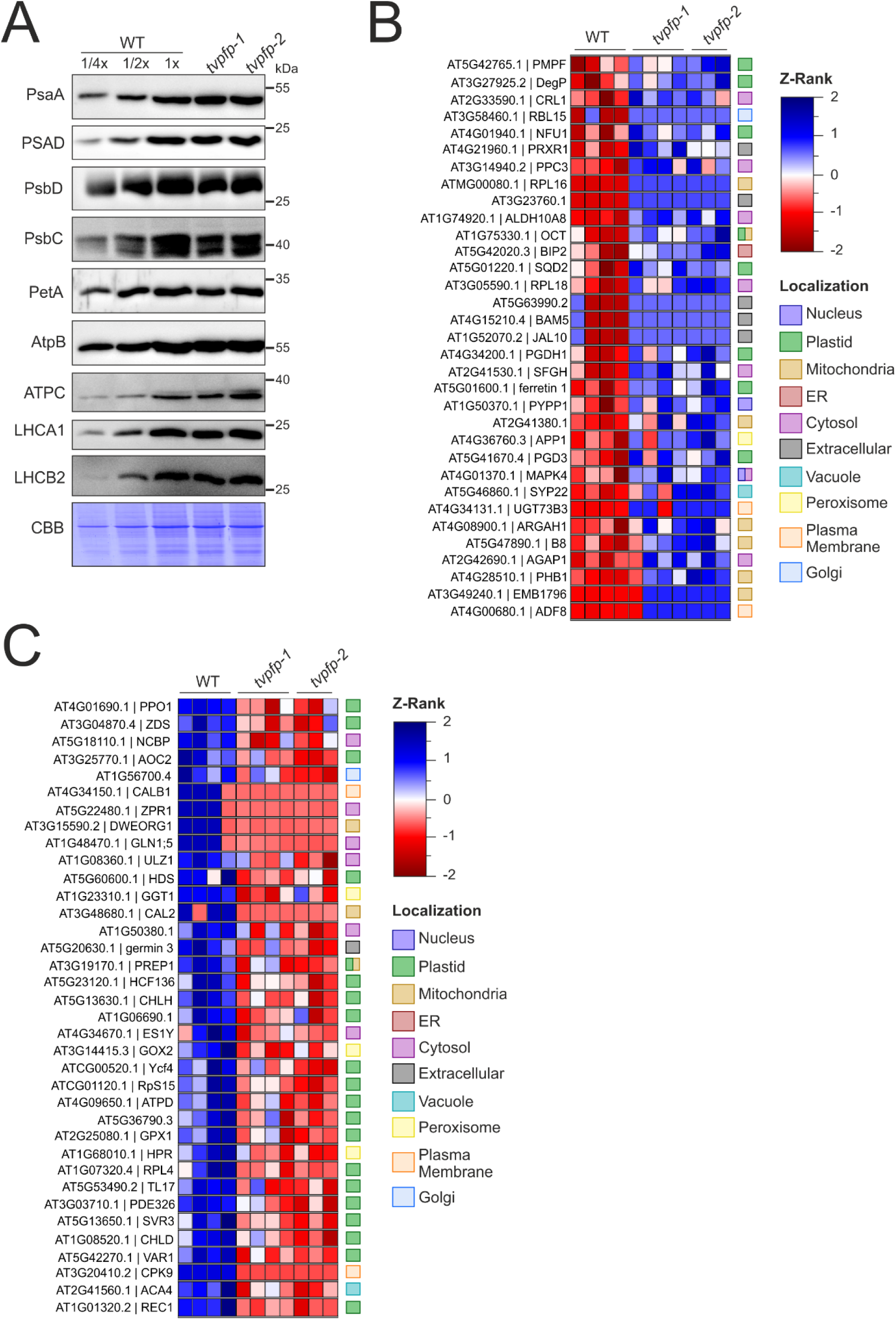
Protein profiles in WT and *tvpfp* mutants. **A)** Accumulation of diagnostic proteins of the major photosynthetic complexes in the wild type (WT) and *tvpfp* mutants. Total protein extracts equivalent to 20 µg were resolved on a 15% PAA gel. The figure shows the accumulation of diagnostic proteins for PSII (PsbD and PsbC) and its associated antenna complex LHCII (LHCB2), PSI (PsaA and PSAD) and its associated antenna complex LHCI (LHCA1), the cytochrome *b*_6_*f* complex (PetA), and the chloroplast ATP synthase (AtpB and ATPC). As a control for equal loading, the Coomassie-stained PAA gel (CBB) is shown below the immunoblot. **B-C)** Total protein profiling by LC-MS/MS. The figure shows proteins **B)** downregulated or **C)** upregulated in *tvpfp-1* and *tvpfp-2* mutants compared with the wild type (WT). The scale indicates Z-rank normalization. N= 3-4 biological replicates; one-way ANOVA, *P* < 0.05 followed Tukeýs HSD post hoc test was employed to identify significant differences in both *tvpfp* mutants compared to WT.

The reduced photosynthetic performance and unchanged accumulation of the major thylakoid-anchored photosynthetic proteins observed in *tvpfp* mutants led us to characterize the proteomes of 12-day-old wild type, *tvpfp-1*, and *tvpfp-2* plants by LC-MS/MS.

Of the 3,636 identified proteins, 470 exhibited statistically significant differential accumulation in at least one sample (, Dataset S1). Among these, 69 proteins were found to be differentially accumulated in both *tvpfp* mutants compared to the wild type, comprising 33 downregulated and 36 upregulated proteins (Fig. 2B, C). Gene ontology (GO) enrichment analysis revealed the upregulation of biological processes associated with photorespiration, pigment biosynthesis (including the tetrapyrrole and carotenoid pathways), and photosynthesis in both *tvpfp* mutants (Fig. S5A). By contrast, wild-type plants showed enrichment of biological processes associated with amino acid metabolism, endoplasmic reticulum stress responses, responses to metal ions and inorganic substances, and catabolic processes (Fig. S5B). Compared with *tvpfp* mutants, wild-type plants also exhibited higher accumulation of the chloroplast envelope-localized protein Plasma Membrane Fusion Protein (PMFP). Together, these results indicate that TVPFP does not play a primary role in the accumulation of major photosynthetic complexes, but influences the accumulation of proteins associated with photorespiration and pigment synthesis. Furthermore, our data suggest that TVPFP affects the accumulation of PMFP (Fig. 2B).

### TVPFP localizes to the chloroplast envelope and undergoes post-translational modification upon relocation to thylakoid membranes

Next, we generated complemented lines expressing a TVPFP-GFP fusion protein under the control of the *UBQ10* promoter (P_ubq10_) in the *tvpfp-1* mutant background (TVPFP-C) (Fig. S6A, B). Semi-quantitative RT-PCR revealed higher transcript levels for *TVPFP* in the complemented lines than in the wild type (Fig. S6C). All complemented lines exhibited wild type-like phenotypes under mixotrophic conditions (Fig. 3A). The accumulation of TVPFP-GFP in total protein extracts from the wild type, *tvpfp-1* and the complemented lines was assessed by PAA gel electrophoresis followed immunodetection confirming the accumulation of TVPFP-GFP in all transgenic lines. However, TVPFP-C2 accumulated lower levels of TVPFP-GFP than the TVPFP-C1 and TVPFP-C3 lines (Fig. 3B).

**Figure 3.**
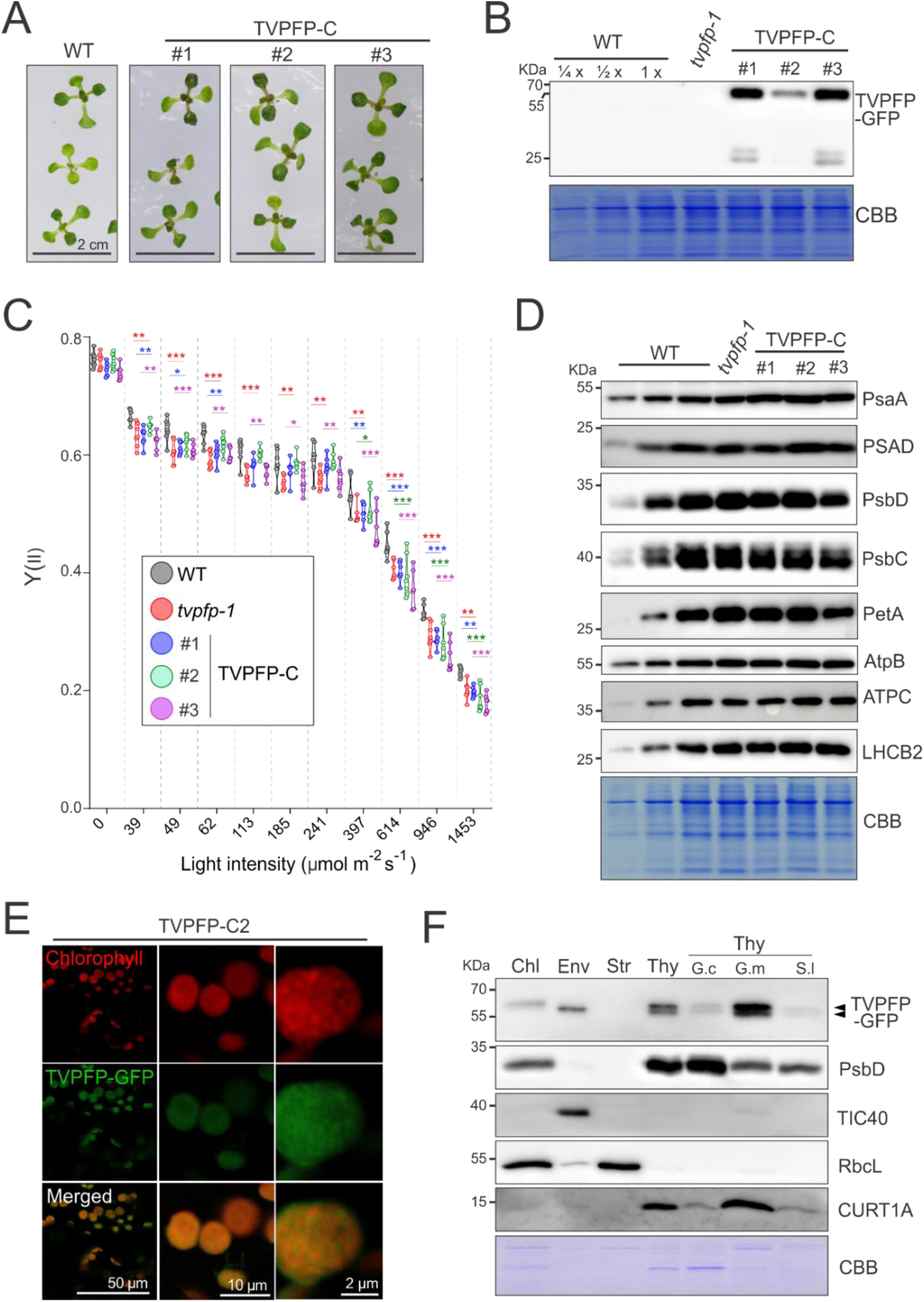
Characterization of TVPFP in TVPFP-C lines. **A)** Phenotype of 12-day-old TVPFP-C lines and wild type (WT) plants grown under mixotrophic conditions. Scale bar = 2 cm. **B)** Immunodetection of TVPFP-GFP in TVPFP-C lines. Total protein extracts (20 µg) were resolved on a 12% PAA gel. The figure shows the accumulation of TVPFP-GFP in TVPFP-C lines 1, 2, and 3. Coomassie Brilliant Blue (CBB) staining was used to confirm equal loading. **C)** Light-response curves of the quantum yield of photosystem II (Y_(II)_) in 12-day-old WT, *tvpfp-1*, and TVPFP-C lines grown under mixotrophic conditions. Statistical differences between the lines and WT were assessed using two-way ANOVA followed by Dunnett’s post hoc test. * *P* < 0.05, \*\**P* < 0.01, \*\*\**P* < 0.001. n = 7 biological replicates. **D)** Accumulation of diagnostic proteins of the major photosynthetic complexes in WT, *tvpfp-1*, and TVPFP-C lines. Total protein extracts (20 µg) were resolved on a 15% PAA gel. The figure shows the accumulation of diagnostic proteins for PSII (PsbD and PsbC) and its associated antenna complex LHCII (LHCB2), PSI (PsaA and PSAD), the cytochrome *b*_6_*f* complex (PetA), and the chloroplast ATP synthase (AtpB and ATPC). As a control for equal loading, the Coomassie-stained PAA gel (CBB) is shown below the immunoblot. **E)** Laser-scanning confocal microscopy of the TVPFP-C2 line. The images show chlorophyll fluorescence (red), TVPFP-GFP fluorescence (green), and the merged image. Three different magnifications are shown. From left to right, the scale bars represent 50, 10, and 2 µm. **F)** Chloroplast fractionation and thylakoid subfractionation of the TVPFP-C2 line. The figure shows protein extracts from chloroplasts (Chl), envelopes (Env), stroma (Str), thylakoid membranes (Thy), and the thylakoid subdomains grana core (G.c), grana margins (G.m), and stroma lamellae (S.l). Fractions equal to 10 µg of protein were resolved on a 15% PAA gel, and immunodetection was performed using diagnostic markers for each fraction, including PsbD (thylakoids and grana core), TIC40 (envelope), RbcL (chloroplasts and stroma), and CURT1A (thylakoids and grana margins). An anti-GFP antibody was used to detect TVPFP-GFP, and the bands corresponding to TVPFP-GFP are indicated by arrows. N = 3 biological replicates.

We also evaluated the photosynthetic performance of 12-day-old TVPFP-C plants grown under mixotrophic conditions using PAM fluorometry, and compared it with that of *tvpfp-1* and wild-type plants (Fig. 3C). The TVPFP-C1 and TVPFP-C3 lines exhibited lower Y_(II)_ values, comparable to those observed in *tvpfp-1* mutants in all light intensities. Compared to the wild type, the TVPFP-C2 line exhibited reduced Y_(II)_ values under high light intensities (397–1453 µmol photons m^-2^ s^-1^) (Fig. 3C). The reduced photosynthetic performance of TVPFP-C lines under high light conditions was largely explained by increased Y(NO) values (Fig. S7A). In addition, the TVPFP-C3 lines exhibited significantly lower Y(NO) values across all tested light intensities (Fig. S7A).

The TVPFP-C lines exhibited NPQ values similar to those of wild-type plants, whereas the *tvpfp-1* mutants displayed increased NPQ values at moderate to high light intensities (113–946 µmol photons m^-2^ s^-1^) (Fig. S7B).

While *tvpfp-1* mutants displayed reduced qL values compared to the wild type under low light intensities (39–62 µmol photons m^-2^ s^-1^), the TVPFP-C2 and TVPFP-C3 lines exhibited lower qL values under high light conditions (614–1453 µmol photons m^-2^ s^-1^) (Fig. S7C). These results suggest that TVPFP-C2 partially restores the photosynthetic capacity of *tvpfp-1* under low light conditions, whereas TVPFP overaccumulation in lines TVPFP-C1 and C3 negatively affects photosynthetic performance.

Next, we analyzed the accumulation of diagnostic proteins for the major photosynthetic complexes in total protein extracts from wild type, *tvpfp-1*, and TVPFP-C complemented lines grown under mixotrophic conditions using PAA gel electrophoresis followed by immunodetection (Fig. 3D). Complemented lines TVPFP-C1 and -C2 accumulated similar contents of protein subunits for PSI (PsaA and PSAD), PSII (PsbD and PsbC) and its associated antenna complex LHCII (LHCB2), the cytochrome *b*_6_*f* complex (PetA), and the chloroplast-localized ATP synthase complex (AtpB and ATPC) (Fig. 3D). However, TVPFP-C3 lines exhibited mild decreased in contents for PSII and the Cyt*b*_6_*f* complex (Fig. 3D).

To avoid artifacts associated with TVPFP overaccumulation, we used the TVPFP-C2 line for further protein characterization. Confocal laser-scanning microscopy revealed chloroplast localization of TVPFP-GFP (Fig. 3E). In addition to the overlap between TVPFP-GFP fluorescence and chlorophyll autofluorescence, TVPFP-GFP signals were also detected at the periphery of chloroplasts, suggesting dual localization of the protein to the envelope and thylakoid membranes (Fig. 3E).

The suborganellar localization of TVPFP was further assessed by chloroplast fractionation and thylakoid sub-fractionation followed by immunodetection (Fig. 3F). Immunoblot analysis indicated that TVPFP was associated with membrane fractions, and was enriched in both envelope and thylakoid membrane (Fig. 3F). Furthermore, thylakoid sub-fractionation revealed enrichment of TVPFP in the grana margin fraction, as well as a putative post-translational modification (PTM) associated with an increase in the molecular weight of TVPFP upon localization to the thylakoid membranes (Fig. 3F).

Next, we analyzed the membrane association strength of TVPFP in thylakoid-enriched samples from TVPFP-C2 plants by salt-wash treatments followed by immunodetection (Fig. S8A). Similar to LHCB2, TVPFP remained in the pellet fraction after sodium chloride and sodium bicarbonate washes (Fig. S8A). In contrast, AtpB was released from the membranes by chaotropic salt treatments (Fig. S8A). We further analyzed the topology of TVPFP by trypsin protection assays in thylakoid membranes isolated from TVPFP-C2 plants. (Fig. S8B). Similar to AtpB, the GFP-tagged C-terminal domain of TVPFP was susceptible to trypsin degradation, whereas PsbO, a lumenal thylakoid protein, exhibited partial resistance to trypsin treatment (Fig. S8B). Together, these results indicate that TVPFP is localized to chloroplast membranes and undergoes post-translational modification upon association with grana margins within the thylakoid membranes. Furthermore, TVPFP is tightly membrane-anchored with the C-terminal domain exposed toward the stromal side.

### TVPFP is present in high-molecular weight complexes

In bacteria, the Tvp38-homologous YqjA proteins are found as monomers, dimers and oligomers (18). Therefore, we explored the putative oligomerization of TVPFP in thylakoid-enriched fractions from the wild type and TVPFP-C2 plants using BS3 chemical crosslinking followed by SDS-PAA electrophoresis. Our results showed that TVPFP accumulates in high-molecular-weight complexes and is detected in both monomeric and oligomeric forms (Fig.4A). To further confirm these findings, we performed blue-native PAGE (BN-PAGE) using thylakoid-enriched fractions from TVPFP-C2 and wild-type plants (Fig. 4B). In-gel fluorescence analysis of the first-dimension PAGE gel revealed three complexes containing TVPFP-GFP, distributed from low to high molecular weight and designated complexes I, II, and III, respectively (Fig. 4B). To confirm that the fluorescence originated exclusively from TVPFP-GFP and not from GFP cleavage products, we performed second-dimension SDS-PAA electrophoresis followed by immunodetection (Fig. 4C). The detection of three distinct TVPFP-GFP-containing complexes in the second-dimension blots confirmed that TVPFP forms high-molecular-weight complexes. While the PTM-modified form of TVPFP accumulated in complexes I and II, the intensity of the unmodified form of TVPFP was increased upon association with the high-molecular-weight complex III (Fig. 4C).

**Figure 4.**
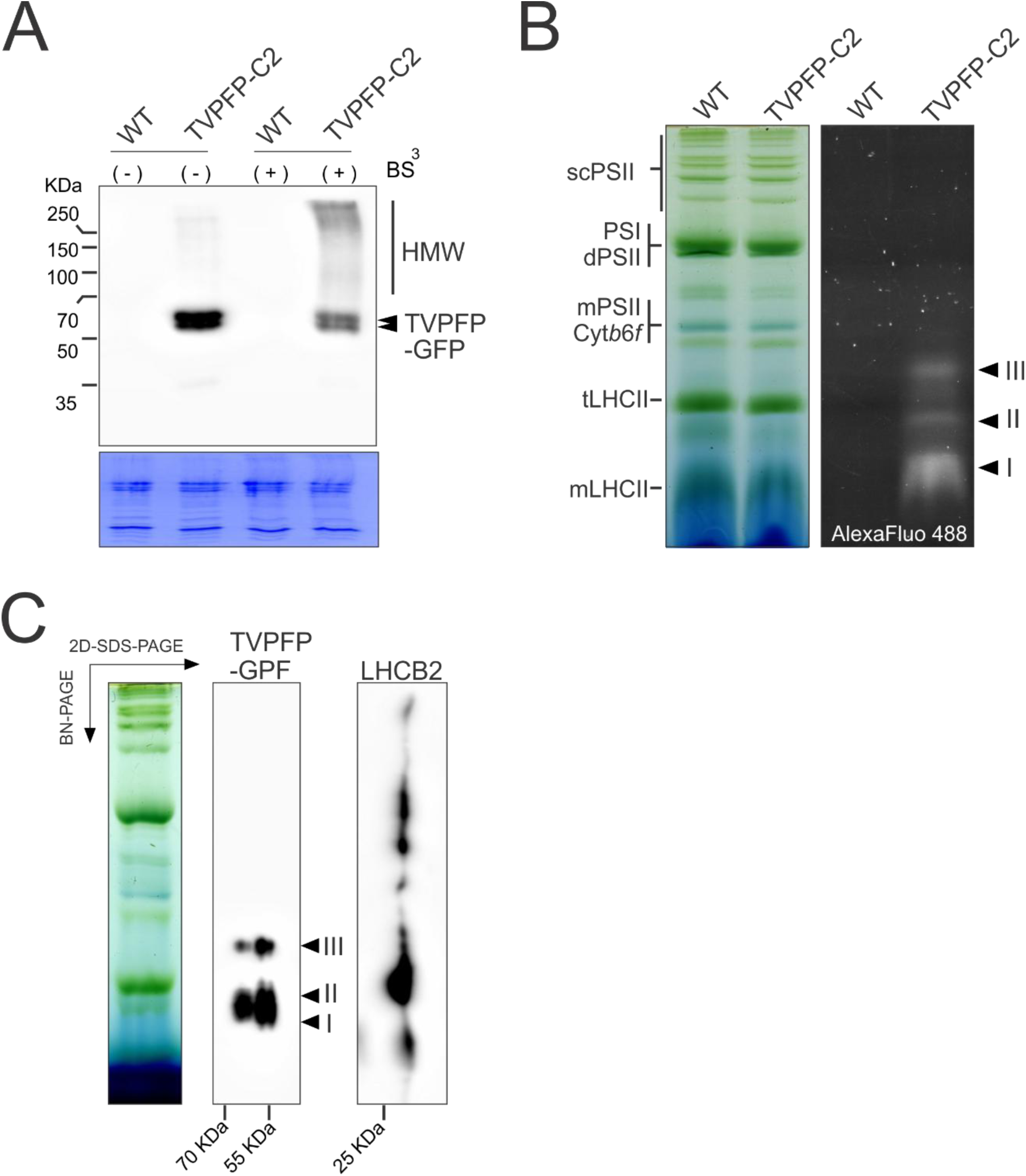
TVPFP-GFP accumulate in high molecular weight complexes. **A)** Chemical cross-linking of thylakoid-enriched membranes from the wild type (WT) and TVPFP-C2 lines. Thylakoid membranes equivalent to 5 µg of protein were incubated with (+) or without (−) BS^3^ cross-linker for 30 min at room temperature and resolved on a 15% SDS-PAA gel. The figure shows the bands corresponding to TVPFP-GFP and the high-molecular-weight (HMW) complexes containing TVPFP-GFP. N = 3 biological replicates. **B)** BN-PAGE (left) followed by fluorescence imaging (right) of thylakoid-enriched fractions from the wild type (WT) and TVPFP-C2 complemented lines. Thylakoid-enriched samples equivalent to 8 µg of chlorophyll were solubilized with the non-ionic detergent β-DDM and resolved on a 4-15% native gradient gel. The main photosynthetic complexes are indicated as PSII supercomplexes (scPSII), PSI and dimeric PSII (PSI, dPSII), monomeric PSII and cytochrome *b*_6_*f* (mPSII, Cyt*b*_6_*f*), LHCII trimers (tLHCII), and monomeric LHCII (mLHCII). For fluorescence imaging, the first-dimension gel was excited at 488 nm to detect GFP fluorescence. GFP-containing complexes were designated complexes I, II, and III and are indicated by black arrowheads. **C)** 2D-SDS-PAGE analyses of thylakoid-enriched fractions from TVPFP-C complemented plants. Thylakoid samples equivalent to 8 µg chlorophyll were solubilized in β-DDM, and resolved in a 4-15% gradient gel. SDS-PAGE was performed as second dimension, and immunodetection was conducted with anti-GFP and anti-LHCB2 antibodies. Putative TVPFP-GFP-containing complexes are indicated in black arrowheads and annotated as I, II, III. N = 3 biological replicates.

### *tvpfp* mutants exhibit reduced vesicle accumulation and decreased abundance of the chloroplast peripheral reticulum

All plant lines investigated exhibited similar chloroplast morphology, with no notable differences between genotypes (Fig. 5A). When we additionally analyzed the grana stack metrics, only slight differences in the key structural parameters were observed, and no clear pattern was associated with the presence of the TVPFP protein. The average grana diameter varied between 466.6 and 541.8 nm (Fig. 5B), while the average grana height was between 136.3 and 165.5 nm, which are typical values for *Arabidopsis thaliana* chloroplasts (Fig. 5C). Additionally, the variability in the values of other grana metrics, including the number of thylakoids building a granum and the grana cross-sectional irregularity (GSI) parameter, was also unrelated to TVPFP accumulation (Fig. 5D, E).

**Figure 5.**
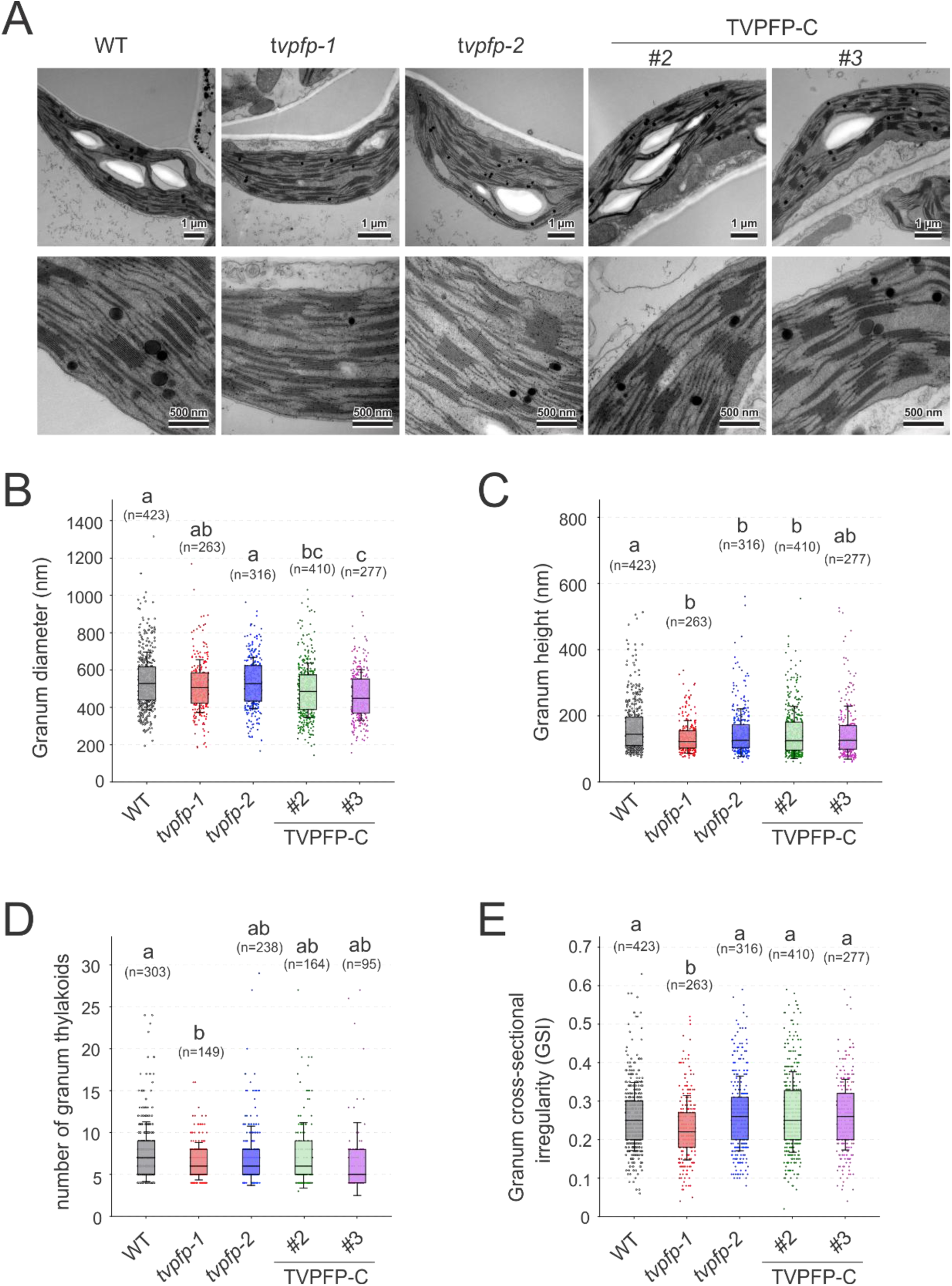
Ultrastructural analysis of the chloroplast thylakoid network and basic grana stack metrics. **A)** TEM micrographs of whole chloroplasts (upper row) and thylakoid network close-ups showing details of grana stack ultrastructure (bottom row) for the wild type (WT), *tvpfp-1*, *tvpfp-2*, TVPFP-C2, and TVPFP-C3 lines. **B-E)** Box plots of selected grana stack metrics extracted using GRANA software. The figure shows metrics for **B)** grana diameter, **C)** grana height, **D)** number of granum thylakoid and **E)** granum cross sectional irregularity (GSI). The box edges show the 25^th^ and 75^th^ percentiles, and error bars indicate SD. Horizontal lines and squares mark the median and mean, respectively. Letters indicate significant differences (*P* < 0.05; Levene’s test for homogeneity of variance, followed by one-way ANOVA with Tukey HSD post hoc test for **b**) and Welch’s ANOVA with Games-Howell post hoc test for **C-E)**; n per variant indicated in the plot legend.

Next, we performed a qualitative analysis of contact sites between the thylakoid network and the inner envelope membrane (Fig. 6). In the analyzed cross-sections, we did not observe any direct connections between the chloroplast envelope and the lamellar thylakoids (Fig. 6A). However, we noticed the presence of the chloroplast peripheral reticulum (CPR), which typically exhibited connections between the inner envelope membrane and the thylakoid stroma or grana membranes within an individual CPR structure (Fig. 6A). It should be noted that, due to the cross-sectional nature of TEM data, these connections were not detected in a single image in all visualized CPR cases.

**Figure 6.**
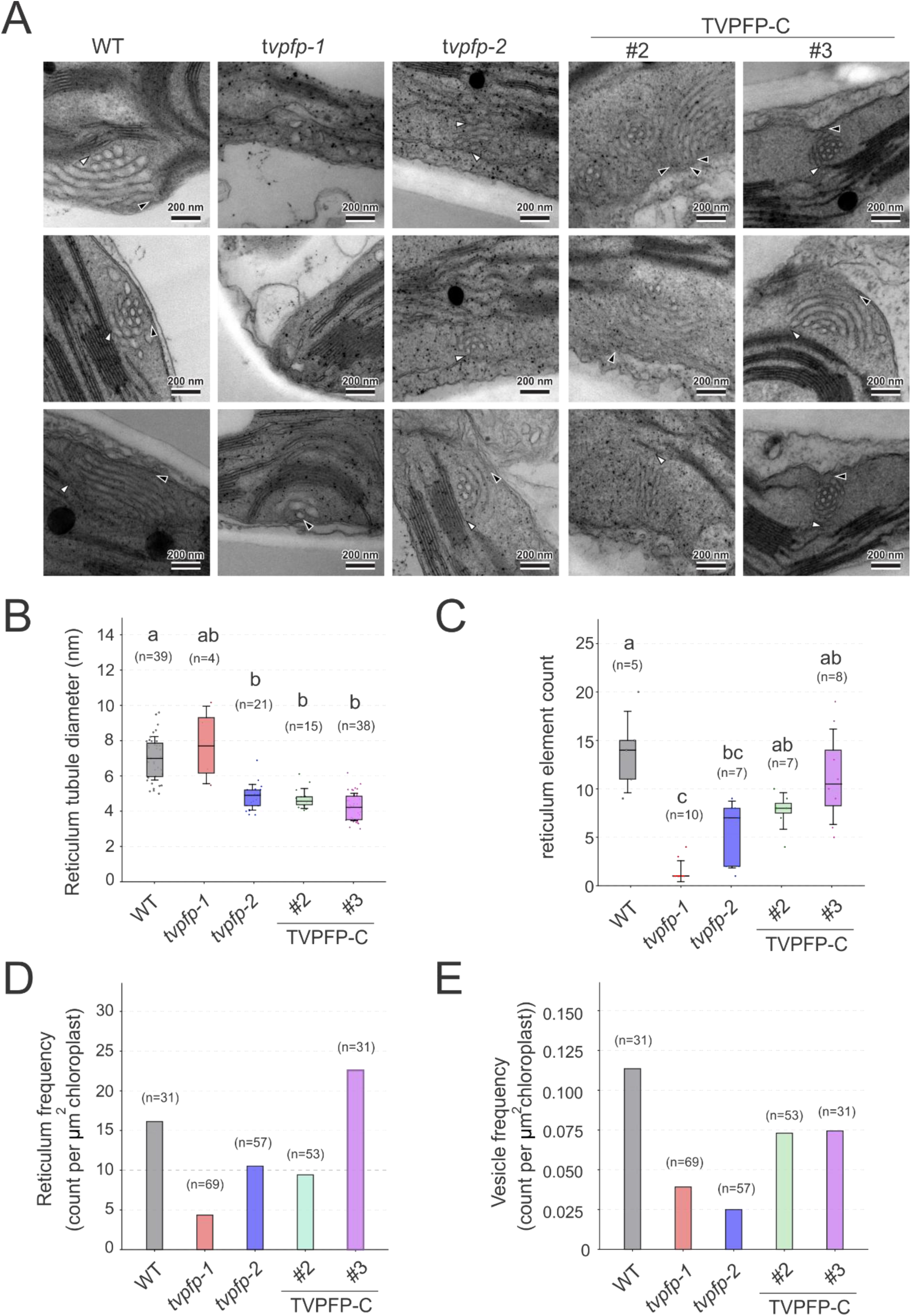
Ultrastructural analysis of the chloroplast peripheral reticulum (CPR). **A)** TEM micrographs of exemplary CPR cross-sections registered in wild type (WT), *tvpfp-1*, *tvpfp-2*, TVPFP-C2, and TVPFP-C3 lines. White arrowheads indicate CPR to thylakoid contact sites. Black arrowheads indicate CPR to inner chloroplast envelope contact sites. **B-C**) Box plot of **B)** reticulum tubule diameter and **C)** reticulum element count. The box edges show the 25^th^ and 75^th^ percentiles, and error bars indicate SD. Horizontal lines and squares mark the median and mean, respectively. Letters indicate significant differences (*P* < 0.05; Levene’s test for homogeneity of variance, followed by Welch’s ANOVA with Games-Howell post hoc test). **D)** Bar chart of CPR frequency per 1 µm² of chloroplast cross-section; n per variant indicated in the plot legend. **E)** Bar chart of vesicle cluster frequency per 1 µm² of chloroplast cross-section. N per variant is indicated in the plot legend.

The CPR of the analyzed Arabidopsis plants is characterized by a specific structure in which tubules with a diameter between 5 and 9.6 nm in wild-type plants (Fig. 6B) are arranged concentrically, and assemblies of 9 to 20 tubules are visible in perpendicular cross-sections (Fig. 6C). Parallel cross-sections reveal that CPR tubules are arranged along the chloroplast envelope and parallel to each other. However, the assemblies also exhibit clear curvature, indicating an arcuate shape of the CPR in *A. thaliana* plants.

Next, we performed a comparative morphometric analysis of the CPR in the wild type, the *tvpfp-1* and *tvpfp-2* mutants, and the TVPFP-C2 and TVPFP-C3 complemented plants. The diameter of the individual tubules building the CPR revealed variability across the lines, without any direct link to the presence of TVPFP (Fig. 6B). However, the analysis of the complexity of the CPR (expressed as the number of elements that make up a single CPR) was reduced in *tvpfp-1* and *tvpfp-2* mutants compared to the wild type, and partially restored in both TVPFP-C lines (Fig. 6C), suggesting that TVPFP is required for the accumulation of CPR, but is not involved in the regulation of CPR tubule size.

We next quantified the abundance of CPR in chloroplast cross-sections (Fig. 6D). The analysis of the frequency of CPR appearance per µm² of chloroplast cross-section showed reduced accumulation of these structures in the *tvpfp-1* and *tvpfp-2* mutants compared to the other lines studied (Fig. 6D). A similar tendency was also evident in the frequency of clustered vesicles appearing in a region similar to CPR, but lacking the structural hallmarks of CPR described above (**Fig. 6e**). Overall, these results suggest an important role of TVPFP in the regulation of CPR abundance and size.

## Discussion

The regulation of processes associated to thylakoid membrane biogenesis and development (known as thylakoistasis) are essential to maintain photosynthesis under changing environmental conditions (19). In this work, we have characterized the role of the only plastid-localized Tvp38/DedA Family Protein in land plants. Tvp38/DedA homologs have been identified from bacteria to higher eukaryotes (20). While yeast Tvp38 homologs play a role in organizing vesicle structures and may contribute to the secretory pathway (21), the thylakoid-less cyanobacterium *Gloeobacter violaceus* PCC 7421 lacks the DedA homolog of the S*ynechocystis* Slr0305 protein, suggesting that DedA may contribute to thylakoid membrane formation in cyanobacteria (20). Most recently, fractionation experiments identified new components of the envelope membrane in a light thylakoid membrane fraction (lightThy) (22). Among the identified proteins, TVPFP and PMPF emerged as candidates mediating communication between the chloroplast envelope and the thylakoids (22). Our results show that the plastid-localized Tvp38 homolog, TVPFP, does not play a role in thylakoid biogenesis or plant development, but is required for fine-tuning photosynthesis.

CPR structures have been described in plants conducting C_3_ or C_4_ photosynthesis, with an inverse correlation between the occurrence of CPR and photorespiration (14, 15). Consistent with these observations, *tvpfp* mutants exhibit lower accumulation of CPR, and high accumulation of photorespiration-related enzymes. In addition, we observed overaccumulation of pigment-related enzymes in *tvpfp* mutants that may represent an attempt to compensate for the low supply of pigments from the envelope membrane to thylakoids, and contribute to the observed similar accumulation levels of protein subunits for the main photosynthetic complexes. In addition, the higher accumulation of the envelope-localized protein PMFP in wild-type plants suggests that TVPFP contributes to the accumulation of PMPF. Thus, the two proteins may act synergistically in the regulation of thylakoid-to-envelope communication.

Similar to the bacterial Tvp38 homolog YqjA (18), TVPFP associates with protein complexes of different molecular weights. Moreover, a PTM reduced the accumulation of TVPFP-containing complexes, suggesting that PTM may contribute to the inactivation of TVPFP upon thylakoid localization. Although TVPFP clearly contributes to vesicle accumulation, the extent of vesicle accumulation was not directly correlated with TVPFP abundance, in contrast to the dosage-dependent phenotype reported for CPSFL1 (8). Instead, our results suggest that TVPFP plays a primary role in the biogenesis of CPR structures in chloroplasts. However, TVPFP does not appear to contribute to the bending of CPR tubules, suggesting that other chloroplast proteins exist that contribute to the maintenance and shaping of these structures.

Overall, our results indicate that TVPFP contributes to the biogenesis of CPR structures and the fine-tuning of photosynthesis, potentially by mediating macromolecular trafficking between the envelope and the thylakoid membranes.

## Materials and Methods

### Plant material and growth conditions

For mixotrophic cultivation, *Arabidopsis thaliana* plants were cultivated in 0.5x MS medium supplemented with 1 % (w/v) sucrose under long day conditions (16 h light, 8 h dark) at 22°C, with light intensity of 120 µmol photons m^-2^ s^-1^. For autotrophic cultivation, *Arabidopsis thaliana* plants were cultivated on soil under long-day conditions (16 h light, 8 h dark), with daytime and nighttime temperatures of 22 °C and 18 °C, respectively, light intensity of 120 µmol photons m^-2^ s^-1^ and 60% humidity. Arabidopsis thaliana seeds were surface sterilized and stratified at 4 °C in the dark for 2-3 days before cultivation. The ecotype Col-0 was used as wild type. The T-DNA insertion mutant *tvpp-1* (SALK_095443) was obtained from NASC under the stock ID N65685. All other transgenic lines, including *tvpp-2* mutants and the complemented lines, were generated in this study.

### Etiolation and de-etiolation kinetics

Etiolation and de-etiolation experiments were conducted according to Sandoval-Ibanez et.al. (2022) (23) with modifications. Briefly, Arabidopsis seeds were surface sterilized, sown on MS media, and stratified for 3 days. The plates were exposed to light for 1 h, then covered with two layers of aluminum foil and cultivated in the dark for 6 days. After 6 days, the seedlings were placed under continuous light (120 µmol photons m^-2^ s^-1^) and plant material was harvested after 0, 6, 12, 24 and 48 h.

### Nucleic acid extraction, cDNA synthesis and RT-PCR

Total DNA was extracted using the cetyltrimethylammonium bromide (CTAB) method. Briefly, plant material was ground in liquid nitrogen, and resuspended in CTAB extraction buffer [2% (w/v) CTAB, 1.4 mM NaCl, 200 mM EDTA pH 8.0, 100 mM Tris-HCl pH 8.0, 100 mM DTT]. The sample was incubated at 65°C for 30 min and centrifuged at 21,000 x *g* for 5 min. The aqueous solution was transferred to a new tube, and mixed with 0.5 volumes of chlorophorm: isoamyl alcohol (24:1). The

sample was vortexed and centrifuged for 30 min at 21,000 x *g* at room temperature. The aqueous phase was transferred into a new tube and mixed with 1 volume of 100 % (v/v) isopropanol followed by centrifugation at room temperature for 30 min at 21,000 x *g*. The pellet containing the genomic DNA was washed with 2 volumes of 70% ethanol, dried at room temperature and resuspended in ultrapure water.

Total RNA extraction was carried out using the NucleoSpin RNA Plant kit (Macherey-Nagel) according to manufacturés recommendations, and quantified by spectroscopic methods. For cDNA synthesis, total RNA equivalent to 0.5 ug was utilized for reverse transcription using the First strand cDNA synthesis kit (Invitrogen). Transcript accumulation was determined by semi-quantitative PCR using a 1:10 diluted cDNA. The diluted cDNA was used for PCR amplification using primers listed in Table S1.

### Genome editing by CRISPR-Cas9

To generate the *tvpfp-2* mutants, we employed the method described by Ruf et.al. (2019) (24). Briefly, a pair of single gRNAs was incorporated into primers pOS445 and pOS446 (Table S1) and used for PCR amplification with the plasmid pHEE2E-TRI as a template. The PCR product was purified using KITNAME, digested with BsaI, and ligated into the binary vector pJF1031 to generate the plasmid pJF1031-TVPFP. *Arabidopsis thaliana* wild type plants were transformed by *Agrobacterium*-mediated transformation, and seeds were selected on MS medium supplemented with 20 mg L^-1^ hygromycin B. Deletion of the targeted sequence was confirmed by genomic DNA extraction, PCR amplification using the primer pair pOS458 and pOS459 (Table S1), followed by DNA sequencing.

### Generation of P_UBQ10_::TVPFP-GFP/*tvpfp-1* complemented lines

The Greengate method was employed to generate the complemented lines (25). Briefly, the sequence of TVPFP was PCR amplified with primers pOS233 and pOS234 (**Table S1**), using the cDNA as template. The purified PCR product was digested with BsaI and ligated into the pGGC000 plasmid. The resulting plasmid, pGGC-TVPFP, was combined with the plasmids pGGA006, pGGB003, pGGD001, pGGE001, pGGF005, and pGGZ001. The resulting mixture was digested with BsaI and ligated using T4 DNA ligase under the following conditions: an initial incubation at 37 °C for 10 min; 28 cycles of 37 °C for 1 min and 15 °C for 3 min; a final incubation at 37 °C for 10 min; and enzyme denaturation at 85 °C for 5 min. The mix was finally transformed into *E.coli* TOP10 and selected for the pGGZ-P_UBQ10_::TVPFP-GFP binary plasmid.

The *tvpfp-1* mutants were transformed by *Agrobacterium*-mediate transformation and the transgenic lines were selected in medium supplemented with 20 mg L^-1^ hygromycin B. The isolation of homozygous P_UBQ10_::TVPFP-GFP lines was conducted by segregation analyses and confirmed by PCR amplification with primers pOS233 and GFP-Rev (**Table S1**).

### Total protein extraction and immunodetection

Total protein extraction was conducted according to Pribil et.al. (2018) (26). Briefly, plant material was snap freezed in liquid nitrogen, ground to a fine powder and resuspended in TKMES buffer [100 mM Tricine-KOH pH 7.5, 10 mM KCl, 1 mM MgCl_2_, 1 mM EDTA, 10% (w/v) sucrose, 0.2% (v/v) Triton X-100, 2× Protease Inhibitor Cocktail for plant cell and tissue extracts (Sigma-Aldrich)], and incubated for 15 min on ice. The samples were centrifuged at 21,000 x *g* for 30 min, the solubilized fraction was collected into a new tube, and total protein extracts were quantified with RotiQuant (ROTH) reagent.

For immunoblot analyses, the samples were resolved in a 12% or 15 % SDS-polyacrylamide (PAA) gels and blotted onto PVDF membranes. All the antibodies in this study are listed in **Table S2**.

### Chlorophyll quantification

Chlorophyll contents were determined spectrophotometrically (27). Thylakoid samples were resuspended in buffered 80 %(v/v) acetone. The samples were then incubated for 30 min at 4°C in darkness, centrifuged at 21,000 x *g* for 20 min, and the absorbance of the supernatant was measured.

### Transmission electron microscopy

Leaves were sampled at the indicated time points. After removing the central vascular bundle, approximately 2 × 2 mm rectangles were excised from the leaf blade and chemically fixed at ambient pressure. Primary fixation was carried out in 2.5% (v/v) glutaraldehyde in 50 mM cacodylate buffer pH 7.4 for 2 h at 23 °C. Samples were then rinsed three times (10 min each) in cacodylate buffer and post-fixed overnight at 4 °C in 2% (w/v) OsO₄ freshly diluted from a 4% (w/v) aqueous stock, in 50 mM cacodylate buffer pH 7.4. Following post-fixation, samples were washed twice (10 min each) each in 50 mM cacodylate buffer pH 7.4 and twice more in MilliQ water. The fixed leaf tissue was dehydrated through a graded acetone series: 10% (v/v) for 10 min, 20% for 10 min, 40% for 15 min, 60% for 30 min and 80% for 40 min, followed by four 40-min changes of 100% acetone and an overnight incubation in 100% acetone. Dehydrated samples were infiltrated with Agar 100 epoxy resin through a graded acetone:resin series [3:1, v/v, for 4 h; 1:1 for 4 h; 1:3 for 20 h], then pure resin for 24 h and, finally, resin containing polymerization accelerator for 2 h. The resin was polymerized for 12 h at 30 °C followed by 48 h at 60 °C.

Ultrathin sections (70 nm) were cut from the polymerized blocks with a Diatome diamond knife on a Leica UCT ultramicrotome and mounted on nickel grids. Images were acquired on a JEM 1400 transmission electron microscope (JEOL, Japan) fitted with an 11-Mpix TEM Morada G2 camera (EMSIS GmbH, Germany).

### Thylakoid membrane morphometrics

Grana nanomorphological parameters were quantified automatically with the GRANA software (28), an AI-based tool that detects grana stacks on calibrated TEM micrographs and returns a suite of grana structural parameters. For each experimental variant, TEM images acquired at identical magnification were processed together in a single batch in fully automatic mode. The automated output was then refined by hybrid-intelligence filtering: granum masks and orientation assignments were inspected visually on the GRANA contact-sheet output, and any incorrectly segmented or misoriented stacks were excluded before downstream analysis.

Two structural parameters were determined for the CPR, measured manually in ImageJ (29). Tubule diameter was obtained by outlining an individual tubule, measuring its cross-sectional perimeter and converting it to the diameter of a circle of equivalent perimeter. The number of tubules (perpendicular to the image plane as well as distinguishable when cut at an angle) forming a single unit of chloroplast peripheral reticulum was counted. Both parameters were measured directly from calibrated TEM micrographs.

### Chloroplast isolation, fractionation and thylakoid sub-fractionation

Chloroplast isolation was conducted according to Trentmann et.al 2020 (30). Briefly, fresh plant material was homogenized in a blender in chloroplast homogenization buffer [0.45 m sorbitol, 20 mm Tricine-KOH, pH 8.4, 10 mm EDTA, 10 mm NaHCO_3_, and 0.1% (w/v) BSA]. The homogenized samples were filtered through two layers of Miracloth®, and centrifuged at 800 x *g* for 10 min at 4°C. The pellet was gently resuspended in homogenization buffer, and loaded into a 40% -80% discontinuous Percoll® gradient. The intact chloroplasts were collected from the 40-80% interphase, and washed with chloroplast resuspension buffer [0.3 m sorbitol, 20 mm Tricine-KOH, pH 7.6, 5 mm MgCl_2_, and 2.5 mm EDTA]. Intact chloroplasts were resuspended in shock buffer

[50 mM HEPES pH 7.5, 5 mM sorbitol, 5 mM MgCl_2_], vortexed, and incubated on ice for 5 min. An aliquot of the disrupted chloroplasts was centrifuged at 13.000 x g for 15 min at 4°C, and the soluble fraction (stroma) was transferred to a new tube. To fractionate the chloroplasts into envelopes and thylakoid, the rest of the intact chloroplasts were loaded in a three-step sucrose gradient [from bottom to top: 0.93 M, 0.6 M, and 0.3 M Sucrose]. After ultra-centrifugation at 70,000g for 1 h at 4°C (with no brake, swing-out rotor SW41-Ti, Beckman), the yellow envelope fraction was collected from the interphase between 0.93M and 0.6M. The envelope fraction was diluted with 3 volumes of ultrapure water and collected by ultracentrifugation at 400,000g for 20 min at 4°C (fix angle roto, TLA120.2, Beckman). The pellet at the bottom of the tube (thylakoid membranes) were gently resuspended in storage buffer [50 mM HEPES pH 7.5, 100 mM sorbitol, 10 mM MgCl_2_] with a soft brush.

The fractionation of thylakoid into grana core, grana margins and stroma lamellae were conducted under dim light at 4°C and according to Koochak et.al. 2019 (31) with modifications. Thylakoid fraction equal to 1 mg L^-1^ of chlorophyll was mixed drop by drop with an equal volume of 2% digitonin solution under continuous stirring. The sample was mixed 5 times in a Dounce homogenizer, and incubated under stirring for 5 min. The insoluble material was removed by centrifugation at 900 *x g* at 4°C for 1 min, and the soluble fraction subjected to ultracentrifugation at 40,000 x g for 30 min at 4°C (fix angle roto TLA120.2, Beckman). The soluble fraction was transferred into a new tube, and the pellet (grana core) was resuspended in storage buffer. The soluble fraction was then subjected to ultracentrifugation at 150,000 *x g* for 1.5 h at 4°C (fix angle roto TLA120.2, Beckman). The loose pellet (grana margins) was collected by pipetting, and the tight pellet (stroma lamellae) was resuspended in storage buffer. All the fractions were quantified with RotiQuant (ROTH) reagent. For immunoblot analyses, the samples were resolved in a 15 % SDS-PAA gels and blotted onto PVDF membranes. All the antibodies in this study are listed in Supplementary table 2.

### Thylakoid-enriched membrane isolation from frozen material

Thylakoid isolation was conducted according to Järvi et. al. (2010) (32) with the modifications from Sandoval-Ibanez et. al. (2022) (23). Plant material was harvested and ground in liquid nitrogen and the powder was resuspended in homogenization buffer [50 mM HEPES pH 7.5, 330 mM sorbitol, 1 mM MgCl_2_, 2 mM EDTA, 5 mM sodium ascorbate and 0.1% (w/v) BSA] at 4 °C under dim light. The homogenized tissue was filtered through two Miracloth layers, and centrifuged at 6000 × *g* for 4 min at 4 °C. The pellet was resuspended in shock buffer [50 mM HEPES pH 7.5, 5 mM sorbitol and 5 mM MgCl_2_], and incubated on ice in darkness for 5 min. The samples were then centrifuged at 6000 × *g* for 4 min at 4 °C, the resulting pellet was washed and resuspended in storage buffer [50 mM HEPES pH 7.5, 100 mM sorbitol, and 10 mM MgCl_2_].

### BS^3^ crosslinking

Thylakoid-enriched samples equal to 5 µg of protein were resuspended in Storage Buffer supplemented with 2× Protease Inhibitor Cocktail for plant cell and tissue extracts (Sigma-Aldrich) and 5 mM BS^3^, and incubated at room temperature for 30 min. The crosslinking reaction was then quenched with Laemmli buffer, the samples were resolved on 15% SDS-PAA gels, and blotted onto PVDF membranes.

### Blue native and second-dimensional PAA gel electrophoresis

Blue native PAA gel electrophoresis (BN-PAGE) was conducted according to Järvi et. al. (2010) (32) with the modifications from Sandoval-Ibanez et. al. (2022) (23). Thylakoid-enriched membranes equal to 0.5 µg of chlorophyll L^-1^ resuspended in 25BTH20G [25 mM BisTris-HCl pH 7.0, 20% (w/v) glycerol and 2× Protease Inhibitor Cocktail for plant cell and tissue extracts (Sigma-Aldrich)] were mixed with an equal volume of 2% (w/v) n-dodecyl-β-maltoside (β-DDM) or 2% digitonin, and incubated for 2 min on ice or 8 min at room temperature and agitation, respectively.

The samples were then centrifuged at 21,000 x *g* for 20 min at 4°C and the soluble fraction was transferred and mixed in a 1:10 volume ratio with Serva Blue G buffer [100 mM BisTris-HCl pH 7.0, 0.5 M ε-aminocaproic acid (ACA), 30% (w/v) sucrose and 50 mg mL^−1^ Serva Blue G] (32). The protein complexes were resolved in 4-15% PAA gradient gels. For the analysis of GFP fluorescence, the first-dimensional BN-PAA gels were placed in a ChemiDoc MP Imaging System (Bio-rad) and visualized with the Alexa488 filter. For the 2D-PAA gel electrophoresis, the strips were solubilized in Laemmli buffer [138 mM Tris-HCL pH 6.8, 6 M urea, 22.2% glycerol, 4.3% w/v SDS and 100 mM DTT] for 1.5 h and resolved by 15% SDS-PAGE. For immunoblot analyses the gels were blotted onto PVDF membranes. All the antibodies in this study are listed in Table S2. For in-gel fluorescence, the polyacrylamide (PAA) gels were scanned using a ChemiDoc Go (Bio-Rad) following excitation at 488 nm.

### Pulse-amplitude modulation (PAM) measurements

For chlorophyll *a* fluorescence analyses, a pool of four 12-day-old plants or leaves from 5-week-old plants were dark-adapted for 15 min prior PAM measurements, using the DUAL-PAM 100 (Heinz Walz GmbH, Effeltrich, Germany). The following settings were used for data collection: Measuring mode = Fluo, Analysis mode= SP-Analisis, Gain = 5, Set Damping = 1ms, Measure Light = 5 and Block Frequency = Auto MF-High. The following program was used for acquisition of light curves: F_v_/F_m_ estimation, pause for 5 s, actinic light set at 6 PAR for 30 s, 16 PAR for 30 s, 29 PAR for 30 s, 80 PAR for 30 s, 152 PAR for 30 s, 208 PAR for 30 s, 364 PAR for 30 s, 581 PAR for 30 s, 913 PAR for 30 s and 1420 PAR for 30 s.

### Confocal laser microscopy

Fluorescence visualization was conducted in a Leica Stellaris 8 Falcon confocal microscope. Confocal microscopy was conducted with a Leica Stellaris 8 Falcon confocal microscope. The excitation/emission settings used were 488/505-530 nm for GFP, 512/520-540 nm for YFP 566/50-600 nm for RFP, and 488/680-720 nm for chlorophyll fluorescence. All images were acquired using sequential scanning settings. The images were combined using the ImageJ software (29), considering the maximum intensity of each stack.

### Topology assay

Thylakoid fraction obtained from frozen material corresponding to 10 µg was incubated with or without 100 ng µL^-1^ trypsin for 20 min on ice prior addition of Laemmli buffer. The proteins were denatured at 85°C for 5 min and resolved by 15% SDS-PAGE. Immunoblot were conducted as described above.

### Salt washes

Thylakoid fractions obtained from frozen material equal to 1 ug were treated with 1M NaCl or 100 mM NaHCO_3_ on ice for 1 hour with agitation every 15 min. The samples were then centrifuged at 16,000 *x g* at 4°C for 1 min and the pellet and soluble fractions were resuspended in Laemmli buffer. The samples were denatured at 80°C for 5 min and resolved by 15% SDS-PAGE. Immunoblot analyses were conducted as described above.

### Mass spectrometry

Total protein extracts equal to 10 µg were resolved in the first 1.5 cm of an SDS-PAA gel. The gel pieces were subjected to in-gel trypsin digestion according to Shevchenko et al. (2006) with modifications (33). Gel pieces were de-stained by de-staining solution [50% (v/v) acetonitrile (ACN), 100 mM NH_4_HCO_3_ pH 8.5] until the blue dye was no longer visible. The gel pieces were dehydrated with 100% (v/v) ACN, rehydrated with reducing solution [10 mM DTT, 100 mM NH_4_HCO_3_ pH 8.5], and incubated at 50 °C for 30 min. Subsequently, the gel pieces were dehydrated with 100% (v/v) ACN, rehydrated with alkylation solution [54 mM iodoacetamide, 100 mM NH_4_HCO_3_ pH 8.5], and incubated in the dark for 30 min. Finally, the gel pieces were dehydrated with 100% (v/v) ACN, rehydrated with protein digestion solution [3ng μL^−1^ Trypsin (Promega Sequencing Grade Modified Trypsin, V5113), 50 mm NH_4_HCO_3_], and incubated overnight at room temperature °C. For each sample, tryptic peptides were extracted from the gel pieces using 50% (v/v) ACN, 0.1% (v/v) TFA, and subsequently with a 100% (v/v) ACN, and collected in a single tube. The tryptic peptides were dried in a vacuum centrifuge at 37 °C. The peptides were finally resuspended in 20 µL of 2% (v/v) ACN, 0.1% (v/v) formic acid (FA).

Tryptic peptides were injected into a Vanquish Neo liquid chromatography system (Thermo Fisher Scientific) and separated using C18 reverse phase chemistry with a pre-column (PepMap Neo Trap-Cartridge from Thermo Fisher Scientific) in line with an Aurora Elite AUR3-15075C18 (IonOpticks). Peptides were eluted via a gradient (50 min for co-IP samples, 90 min for total cell extract samples) increasing from 4 to 40% acetonitrile in 0.1% formic acid and electrosprayed via Nanospray Flex ion source into an Exploris 480 mass spectrometer (Thermo Fisher Scientific). Samples were measured with a data-dependent acquisition scan strategy with one MS full scan with a resolution of 120,000 followed by TopSpeed MS/MS fragment ion scans with a cycle time of 2 s and a resolution of 15,000.

MS raw files were analyzed using MaxQuant, version 2.5.1.0 and the Andromeda algorithm (34, 35). The search engine was used with default settings, including PSM, peptide and protein FDR set at 1% and match between runs enabled. The *A. thaliana* database (Araport11; https://www.arabidopsis.org) with contaminants included was used as the analytical basis. Precursor mass tolerance was set to 20 ppm and the fragment ion tolerance was 0.5 Da. Carbamidomethylation of cysteines was set as the fixed modification and oxidation of methionine, acetylation of protein N-termini, pyroglutamate formation, phosphorylation of serine and threonine and in case of the co-IP samples thioacyl as variable modification, with two missed cleavages allowed. For quantification by the MaxLFQ algorithm (36) a minimum ratio count of two unique or razor peptides was required. The MS data were uploaded to the ProteomeXchange consortium at PRIDE (https://www.ebi.ac.uk/pride) and are available under accession PXDxxxxxx.

Perseus software was used for LFQ analysis (37, 38). Contaminants were removed, datasets corresponding to each genotype were grouped, and the data were log2-transformed. Proteins with valid values in less than 70% of the samples in at least one genotype were removed, and missing values were imputed as zero. The data were subjected to an ANOVA multiple-sample test with a significance threshold of *P* < 0.05. Proteins showing significant differences in at least one comparison were retained and subjected to a post hoc Tukey’s HSD test with an FDR < 0.05. Protein pairs showing significant differences in both *tvpfp* mutants compared with the wild type were considered true differentially expressed proteins associated with TVPFP. Gene ontology enrichment was determined with ShinyGO tool using default settings (39).

## Acknowledgments

We thank the confocal microscopy facility of Ruhr University Bochum (DFG Project no. 523980288, GZ: INST 213/1180-1 FUGG). TEM studies were performed in the Laboratory of Electron Microscopy of the Nencki Institute of Experimental Biology, supported by a project financed by the Polish Minister of Education and Science under contract no. 2022/WK/05 (Polish Euro-BioImaging Node “Advanced Light Microscopy Node Poland”). O.S-I thanks Dr. Stéphanie Arrivault for critical reading of the manuscript.

## References

1. M. P. Johnson, Photosynthesis. Essays Biochem 60, 255–273 (2016).

2. D. Rolo et al., CO-EXPRESSED WITH PSI ASSEMBLY1 (CEPA1) is a photosystem I assembly factor in Arabidopsis. Plant Cell 36, 4179–4211 (2024).

3. J. Komenda, R. Sobotka, P. J. Nixon, The biogenesis and maintenance of PSII: Recent advances and current challenges. Plant Cell 36, 3997–4013 (2024).

4. A. A. Kelly et al., Synthesis and transfer of galactolipids in the chloroplast envelope membranes of Arabidopsis thaliana. Proc Natl Acad Sci U S A 113, 10714–10719 (2016).

5. M. Ostermeier, A. Garibay-Hernandez, V. J. C. Holzer, M. Schroda, J. Nickelsen, Structure, biogenesis, and evolution of thylakoid membranes. Plant Cell 36, 4014–4035 (2024).

6. E. Lindquist, H. Aronsson, Chloroplast vesicle transport. Photosynth Res 138, 361–371 (2018).

7. N. Z. Khan, E. Lindquist, H. Aronsson, New putative chloroplast vesicle transport components and cargo proteins revealed using a bioinformatics approach: an Arabidopsis model. PLoS One 8, e59898 (2013).

8. A. P. Hertle et al., A Sec14 domain protein is required for photoautotrophic growth and chloroplast vesicle formation in Arabidopsis thaliana. Proc Natl Acad Sci U S A 117, 9101–9111 (2020).

9. W. Wietrzynski et al., Molecular architecture of thylakoid membranes within intact spinach chloroplasts. Elife 14 (2025).

10. B. D. Engel et al., Native architecture of the Chlamydomonas chloroplast revealed by in situ cryo-electron tomography. Elife 4 (2015).

11. A. Rast et al., Biogenic regions of cyanobacterial thylakoids form contact sites with the plasma membrane. Nat Plants 5, 436–446 (2019).

12. T. Armarego-Marriott, O. Sandoval-Ibanez, L. Kowalewska, Beyond the darkness: recent lessons from etiolation and de-etiolation studies. J Exp Bot 71, 1215–1225 (2020).

13. T. Armarego-Marriott et al., Highly Resolved Systems Biology to Dissect the Etioplast-to-Chloroplast Transition in Tobacco Leaves. Plant Physiol 180, 654–681 (2019).

14. J. H. Hilliard, S. H. West, The association of chloroplast peripheral reticulum with low photorespiration rates in a photorespiring plant species. Planta 99, 352–356 (1971).

15. V. E. Gracen, Jr., J. H. Hilliard, R. H. Brown, S. H. West, Peripheral reticulum in chloroplasts of plants differing in CO2 fixation pathways and photorespiration. Planta 107, 189–204 (1972).

16. S. J. Karpowicz, S. E. Prochnik, A. R. Grossman, S. S. Merchant, The GreenCut2 resource, a phylogenomically derived inventory of proteins specific to the plant lineage. J Biol Chem 286, 21427–21439 (2011).

17. C. M. Hooper, I. R. Castleden, S. K. Tanz, N. Aryamanesh, A. H. Millar, SUBA4: the interactive data analysis centre for Arabidopsis subcellular protein locations. Nucleic Acids Res 45, D1064–D1074 (2017).

18. R. Keller, N. Schleppi, J. Weikum, D. Schneider, Mutational analyses of YqjA, a Tvp38/DedA protein of E. coli. FEBS Lett 589, 842–848 (2015).

19. W. Sakamoto, Thylakostasis: key factors in thylakoid membrane organization with emphasis on biogenesis and remodeling proteins in vascular plants. Plant Cell Physiol 66, 1602–1618 (2025).

20. R. Keller, D. Schneider, Homologs of the yeast Tvp38 vesicle-associated protein are conserved in chloroplasts and cyanobacteria. Front Plant Sci 4, 467 (2013).

21. Y. Hama, H. Morishita, N. Mizushima, Regulation of ER-derived membrane dynamics by the DedA domain-containing proteins VMP1 and TMEM41B. EMBO Rep 23, e53894 (2022).

22. E. W. LaBrant et al., Lipid metabolism and contact site homologs are present at the chloroplast envelope-thylakoid interface. Nat Commun 17 (2026).

23. O. Sandoval-Ibanez et al., De-etiolation-induced protein 1 (DEIP1) mediates assembly of the cytochrome b(6)f complex in Arabidopsis. Nat Commun 13, 4045 (2022).

24. S. Ruf et al., High-efficiency generation of fertile transplastomic Arabidopsis plants. Nat Plants 5, 282–289 (2019).

25. A. Lampropoulos et al., GreenGate---a novel, versatile, and efficient cloning system for plant transgenesis. PLoS One 8, e83043 (2013).

26. M. Pribil et al., Fine-Tuning of Photosynthesis Requires CURVATURE THYLAKOID1-Mediated Thylakoid Plasticity. Plant Physiol 176, 2351–2364 (2018).

27. R. J. Porra, W. A. Thompson, P. E. Kriedemann, Determination of accurate extinction coefficients and simultaneous equations for assaying chlorophylls a and b extracted with four different solvents: verification of the concentration of chlorophyll standards by atomic absorption spectroscopy. Biochimica et Biophysica Acta (BBA) -Bioenergetics 975, 384– 394 (1989).

28. A. Bukat et al., GRANA: An AI-based tool for accelerating chloroplast grana nanomorphology analysis using hybrid intelligence. Plant Physiol 198 (2025).

29. C. A. Schneider, W. S. Rasband, K. W. Eliceiri, NIH Image to ImageJ: 25 years of image analysis. Nat Methods 9, 671–675 (2012).

30. O. Trentmann et al., Identification of Chloroplast Envelope Proteins with Critical Importance for Cold Acclimation. Plant Physiol 182, 1239–1255 (2020).

31. H. Koochak, S. Puthiyaveetil, D. L. Mullendore, M. Li, H. Kirchhoff, The structural and functional domains of plant thylakoid membranes. Plant J 97, 412–429 (2019).

32. S. Jarvi, M. Suorsa, V. Paakkarinen, E. M. Aro, Optimized native gel systems for separation of thylakoid protein complexes: novel super- and mega-complexes. Biochem J 439, 207–214 (2011).

33. A. Shevchenko, H. Tomas, J. Havlis, J. V. Olsen, M. Mann, In-gel digestion for mass spectrometric characterization of proteins and proteomes. Nat Protoc 1, 2856–2860 (2006).

34. J. Cox et al., Andromeda: a peptide search engine integrated into the MaxQuant environment. J Proteome Res 10, 1794–1805 (2011).

35. J. Cox, M. Mann, MaxQuant enables high peptide identification rates, individualized p.p.b.-range mass accuracies and proteome-wide protein quantification. Nat Biotechnol 26, 1367–1372 (2008).

36. J. Cox et al., Accurate proteome-wide label-free quantification by delayed normalization and maximal peptide ratio extraction, termed MaxLFQ. Mol Cell Proteomics 13, 2513– 2526 (2014).

37. J. D. Rudolph, J. Cox, A Network Module for the Perseus Software for Computational Proteomics Facilitates Proteome Interaction Graph Analysis. J Proteome Res 18, 2052– 2064 (2019).

38. S. Tyanova et al., The Perseus computational platform for comprehensive analysis of (prote)omics data. Nat Methods 13, 731–740 (2016).

39. S. X. Ge, D. Jung, R. Yao, ShinyGO: a graphical gene-set enrichment tool for animals and plants. Bioinformatics 36, 2628–2629 (2020).

40. J. Waese et al., ePlant: Visualizing and Exploring Multiple Levels of Data for Hypothesis Generation in Plant Biology. Plant Cell 29, 1806–1821 (2017).

