## Supplementary Information for "Tvp38/DedA Family Protein mediates biogenesis of chloroplast peripheral reticulum and envelope-to-thylakoid contacts in *Arabidopsis*"

**Supplementary Figures**

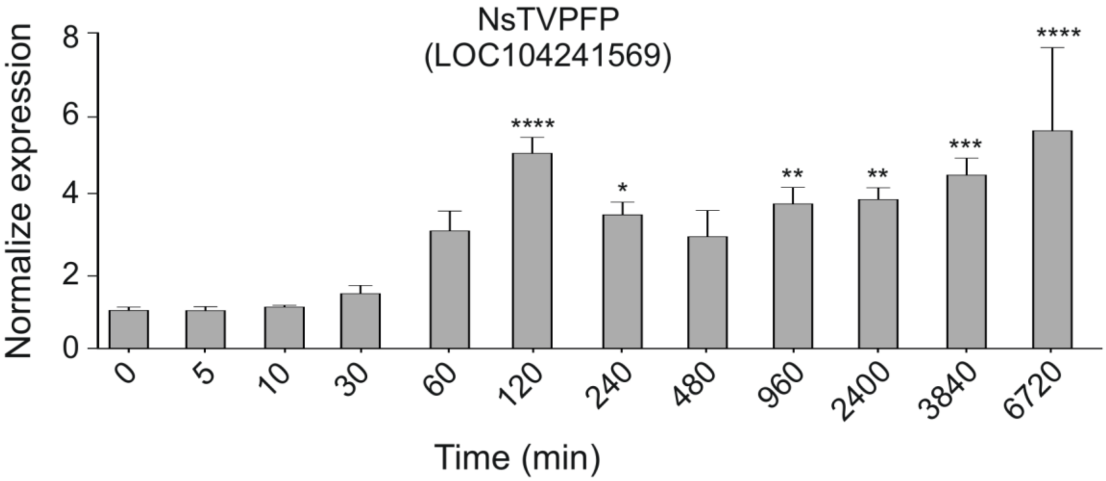


**Fig. S1. Expression pattern of TVPFP in tobacco during de-etiolation.**

Expression of tobacco TVPFP in previously reported time-resolved de-etiolation experiments. The x-axis gives the time of illumination from 0 (T0) to 6720 (T6720) minutes. The y-axis shows DEIP1 transcript accumulation in RPKM. Error bars represent standard deviation; n=3 biological replicates. Statistical significance was determined by one-way ANOVA followed Dunnet post-test. * $P<0.05$, ** $P<0.005$, *** $P<0.001$.

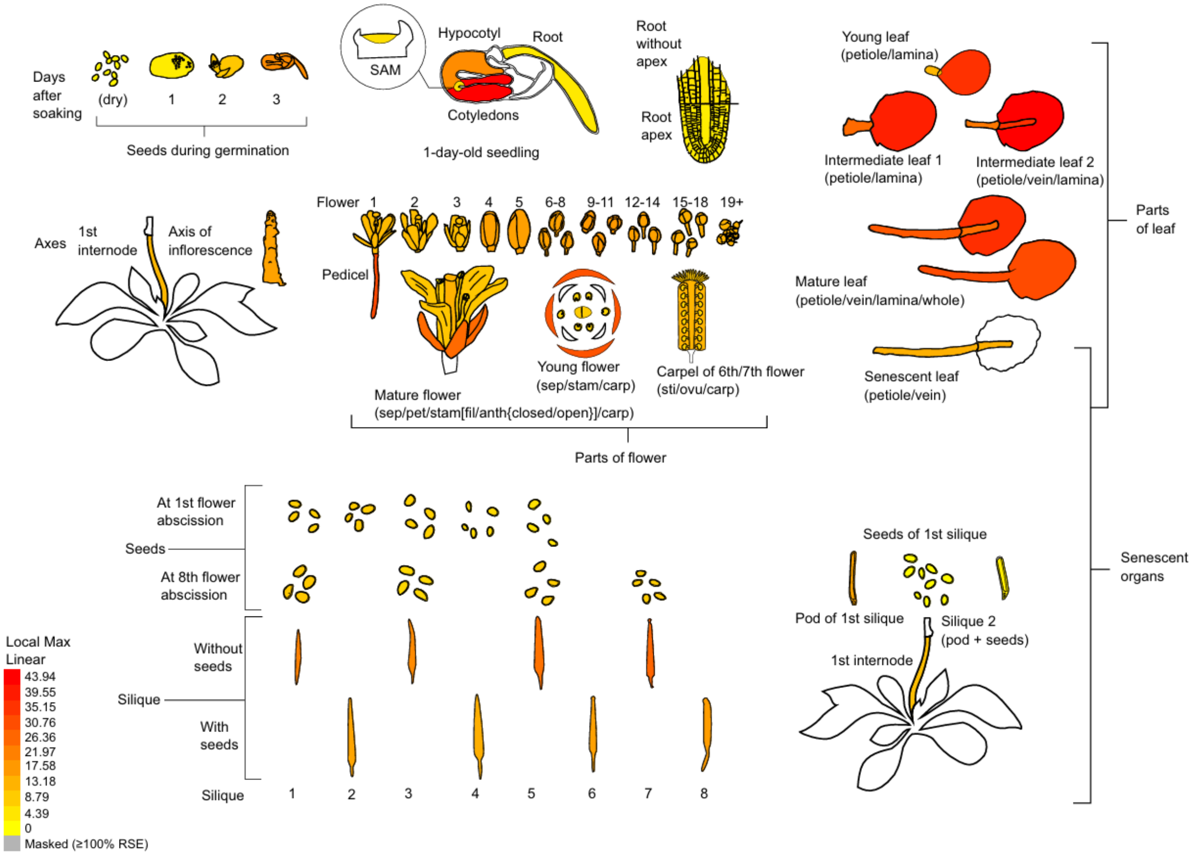


**Fig. S2. Expression of *TVPFP* in Arabidopsis.**

The expression data for AT1G22850 were obtained from the eFP browser (bar.utoronto.ca)(1).

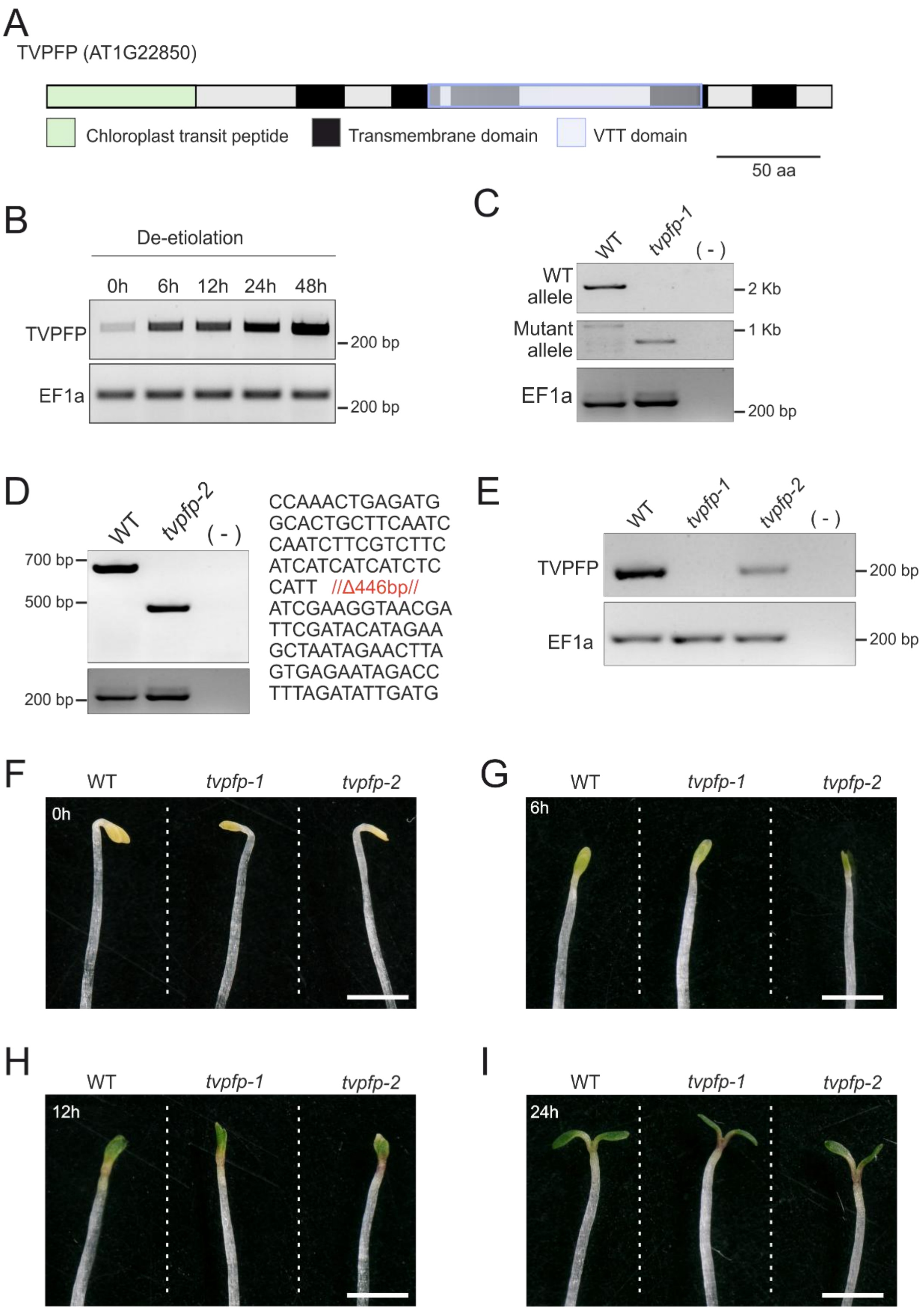


**Fig. S3. Expression of TVPFP during de-etiolation, mutant isolation and plant phenotypes during de-etiolation.**

**A)** Schematic representation of the TVPFP protein. The figure shows the putative chloroplast transit peptide (green), the predicted transmembrane segments (black), and the VTT domain (blue). The scale bar represents 50 amino acids (aa). **B)** Expression of *TVPFP* during de-etiolation determined by semi-quantitative PCR. The figure shows semi-quantitative PCR amplification after 30 cycles for *TVPFP* in etiolated plants (0 h) and after 6, 12, 24, and 48 h of illumination. Amplification of the transcript encoding *EF1a* was used as a control and performed for 28 cycles. n = 3 independent biological replicates. **C)** Genotyping of the *tvpfp-1* insertional mutant line. The left border of the T-DNA was used as the target sequence for amplification of the mutant allele. n = 3 independent biological replicates. **D)** Genotyping of *tvpfp-2*. The genomic sequence of *TVPFP* was amplified by PCR. The figure shows the 446-bp deleted segment in the *tvpfp-2* mutant line. *n* = 3 independent biological replicates. **E)** Expression of *TVPFP* in WT, *tvpfp-1*, and *tvpfp-2*. The figure shows semi-quantitative PCR amplification after 32 cycles for *TVPFP*. Amplification of the transcript encoding *EF1α* was used as a control and performed for 28 cycles. *n* = 3 independent biological replicates. **F-I)** Phenotypes of WT, *tvpfp-1*, and *tvpfp-2* during etiolation and de-etiolation. The figures show **F)** etiolated plants (0 h) and plants after illumination for **G)** 6 h, **H)** 12 h, and **I)** 24 h. n = 3 independent biological replicates.

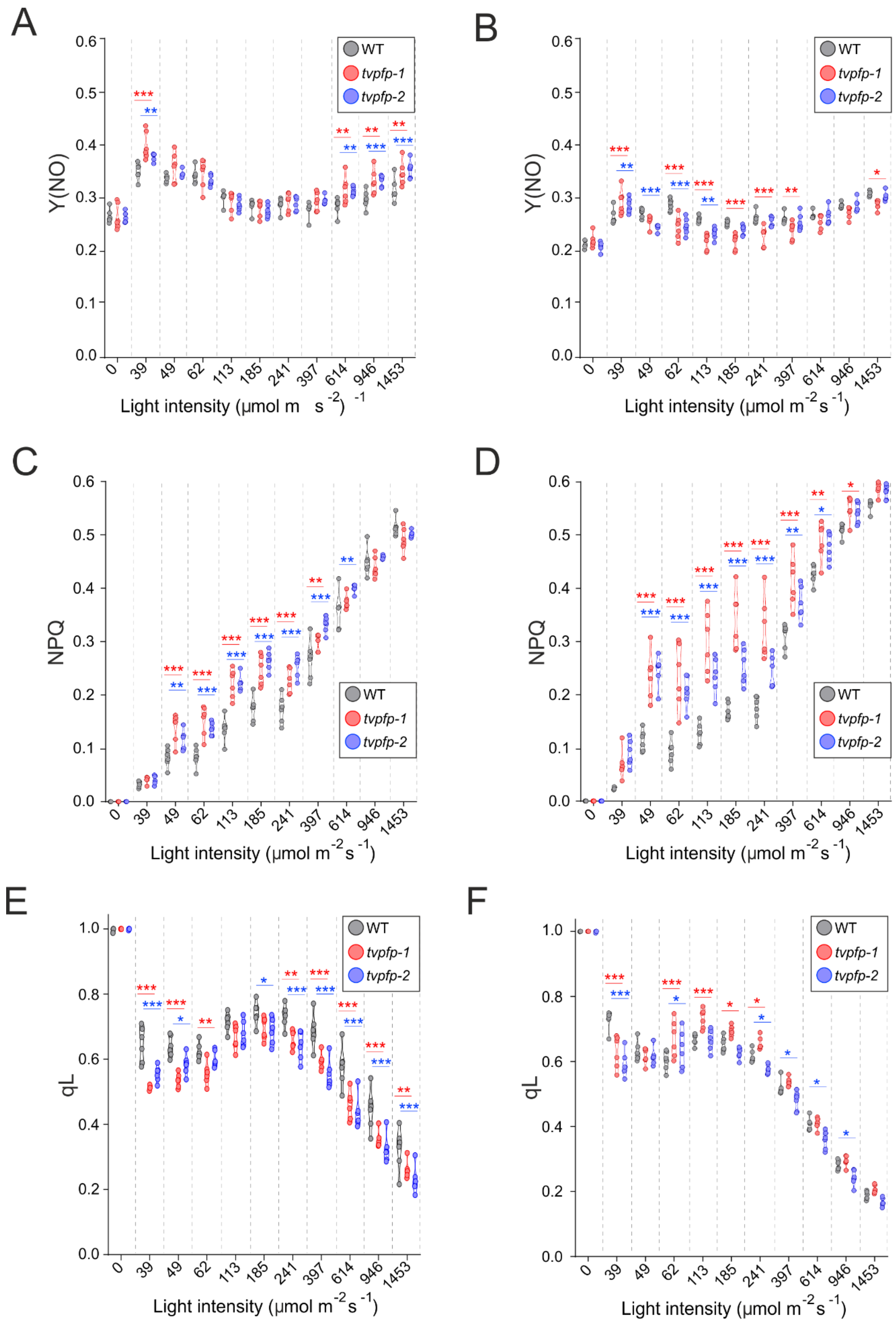
A
B
C
D
E
F
Y(NO)
NPQ
qL
WT
tvpfp-1
tvpfp-2
0
39
49
62
113
185
241
397
614
946
1453
Light intensity (μmol m s -2) -1
Light intensity (μmol m-2 s-1)

**Fig. S4. The chlorophyll *a* fluorescence parameters in WT, *tvpfp-1* and *tvpfp-2*.**

Light-response curves quantum yield of **A-B)** non-Regulated energy dissipation in photosystem II (Y(NO)), **C-D)** non photochemical quenching (NPQ) and **E-F)** open reaction centers of photosystem II (qL) in WT (grey), tvpfp-1 (red), and tvpfp-2 (blue) plants grown under **A, C, E)** mixotrophic conditions or **B, D, F)** photoautotrophic conditions. Statistical differences between the mutants and WT were assessed using two-way ANOVA followed by Dunnett's post hoc test. * $P < 0.05$, ** $P < 0.01$, *** $P < 0.001$. n = 6 biological replicates.

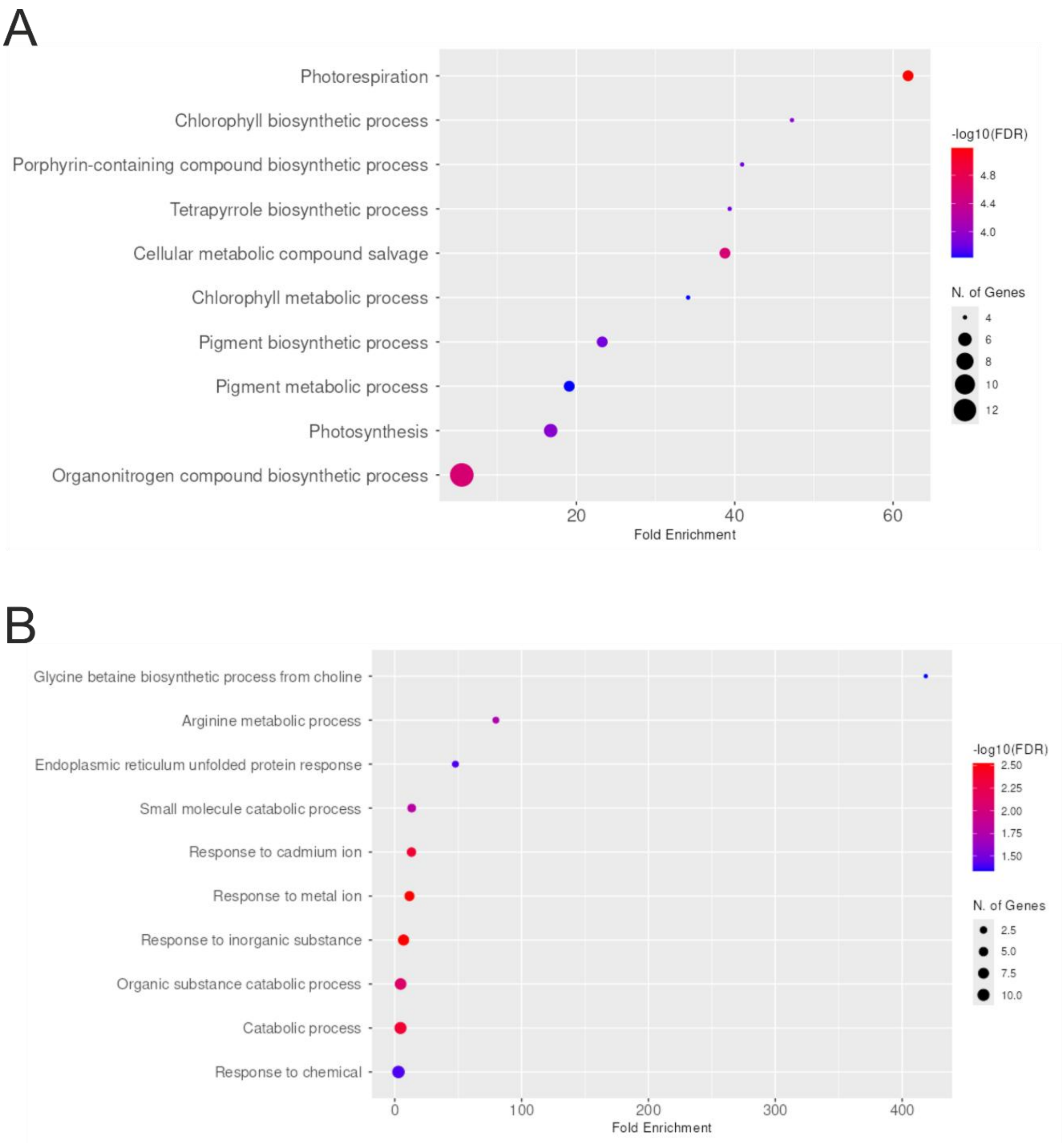


**Fig. S5. Gene Ontology (GO) enrichment analysis for biological processes.**

The figure shows the GO enrichment analysis for **A)** up-regulated and **B)** down-regulated proteins in *tvpfp-1* and *tvpfp-2* mutants compared with the wild type.

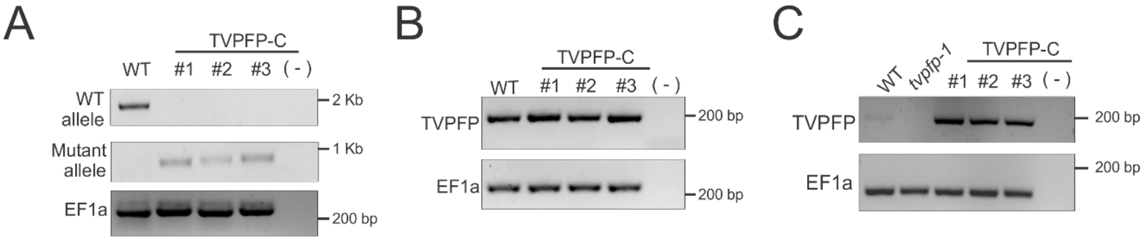


**Fig. S6. Construction of TVPFP-C complemented lines.**

**A)** Genotyping of the *tvpfp-1* mutant background from homozygous TVPFP-C lines. The figure shows the WT and mutant alleles of the *tvpfp-1* mutant. The left border of the T-DNA was used as the target sequence for amplification of the mutant allele. n = 3 independent biological replicates. **B)** Genotyping of TVPFP-GFP. The figure shows the amplification of TVPFP in WT and the TVPFP-C complemented lines. **C)** Expression of TVPFP in WT, *tvpfp-1*, and TVPFP-C complemented lines. The figure shows semi-quantitative PCR amplification after 28 cycles for TVPFP. Amplification of the transcript encoding EF1α was used as a control and performed for 28 cycles. n = 3 independent biological replicates.

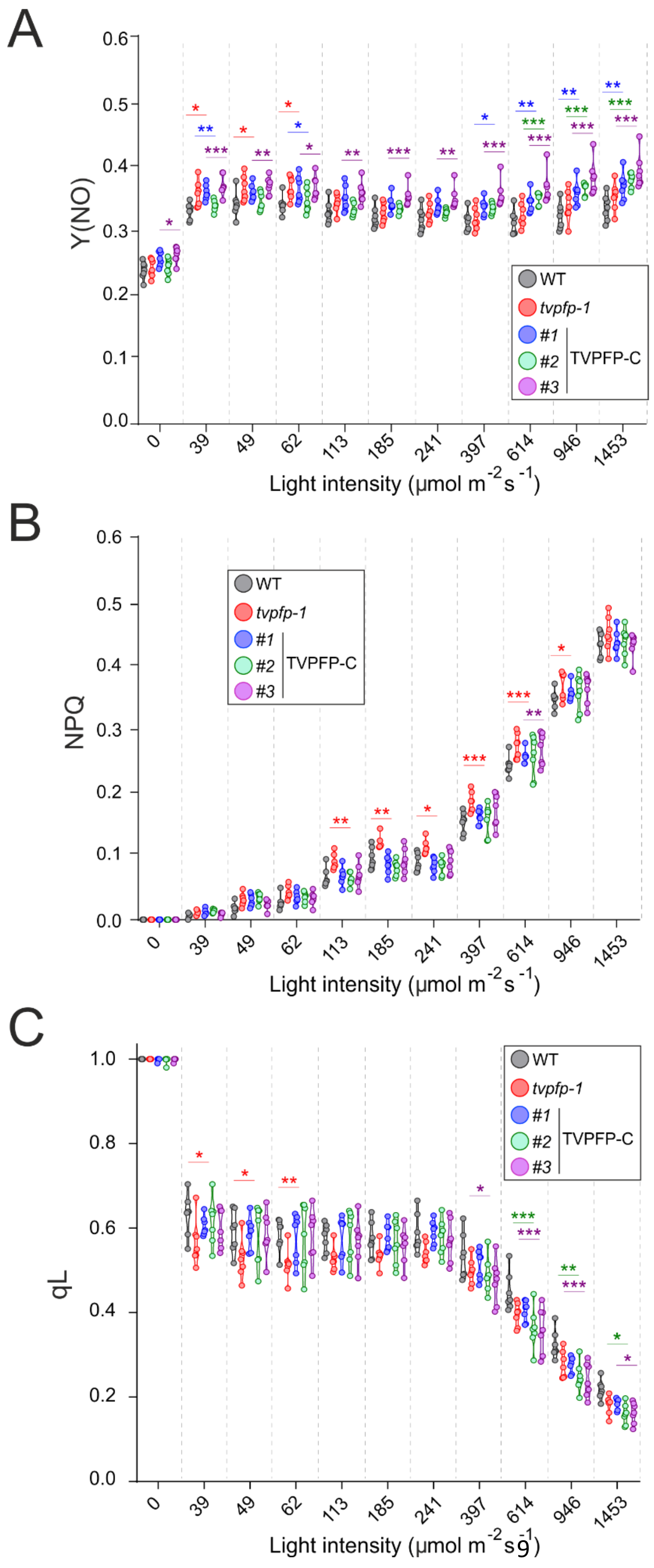
A
Y(NO)
WT
tvpfp-1
#1
#2
#3
TVPFP-C
0.0
0.1
0.2
0.3
0.4
0.5
0.6
0
39
49
62
113
185
241
397
614
946
1453
Light intensity (µmol m-2 s-1)
B
NPQ
C
qL
0.8
1.0

**Fig. S7. The chlorophyll a fluorescence parameter in WT, *tvpfp-1*, and TVPFP-C lines.**

Light-response curves of **A)** quantum yield of non-regulated energy dissipation in photosystem II (Y(NO)), **B)** non photochemical quenching (NPQ) and **C)** open reaction centers of photosystem II (qL) in WT (grey), *tvpfp-1* (red), and TVPFP complemented lines C1 (blue), C2 (green) and C3 (magenta) plants grown under mixotrophic conditions. Statistical differences between the mutants and WT were assessed using two-way ANOVA followed by Dunnett's post hoc test. * $P < 0.05$, ** $P < 0.01$, *** $P < 0.001$. n = 7 biological replicates.

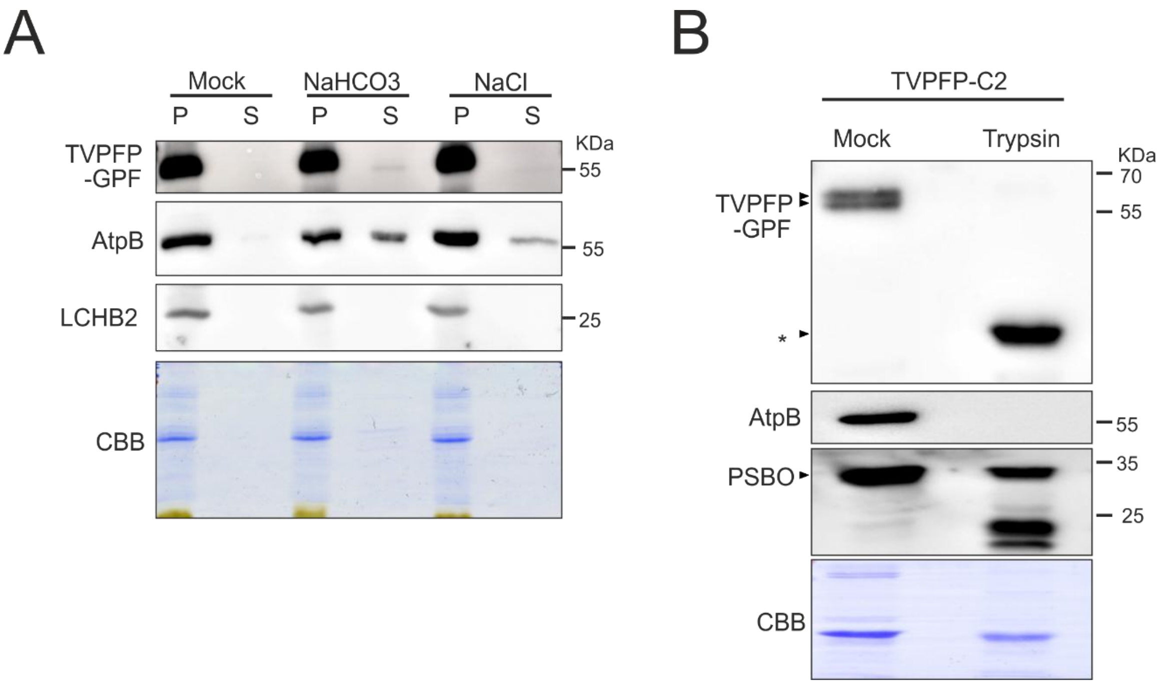


**Fig. S8. The chlorophyll a fluorescence parameter in WT, *tvpfp-1*, and TVPFP-C lines.**

**A)** Salt wash of thylakoid-enriched samples from TVPFP-C2 lines. Thylakoid-enriched samples equal to 1 µg of chlorophyll were incubated with 0.1 M NaHCO3 or 1 M NaCl for 1 hour and separated in pellet (P) and soluble (S) fractions by centrifugation. Proteins present in the pellet and soluble fractions were resolved in a 12% PAA gel, and probed with anti-GFP, anti-AtpB, and anti-LHCB2 antibodies. Coomassie-stained PAA gel (CBB) is shown below the immunoblot. n = 3 independent biological replicates. **B)** Trypsin protection assay. Thylakoid-enriched fractions from TVPFP-C2 lines equivalent to 2 µg of chlorophyll were incubated with 1 µg $mL^{-1}$ trypsin at 25 °C for 30 min, resolved in a 12.5% PAA gel, and probed with anti-GFP for detection of TVPFP-GFP, anti-AtpB antibodies (recognizing a protein containing domains exposed to the stromal side and, therefore, being susceptible to tryptic digestion) and anti-PSBO (detecting a marker protein that resides at the lumenal side of the membrane and, therefore, is protected from tryptic digestion). The TVPFP-GFP degradation products resulting from tryptic digestion are indicated with an asterisk. As a control for equal loading, the Coomassie-stained PAA gel (CBB) is shown below the immunoblot.

**Supplementary Tables**

**Table S1.** Primers used in this study.

| Primer name | Sequence (5´to 3´) | Description |
|---|---|---|
| pOS233 | AACAGGTCTCAGGCTCGATGCGCAGCCTCACTCTAG | Cloning the coding sequence of AT1G22850 and genotyping of *tvpfp-1* mutants. Sequence in red indicate adaptors for Greengate cloning |
| pOS234 | AACAGGTCTCACTGACTTTGCATCTCTCTTCTCATCATC | Cloning the coding sequence of AT1G22850 and genotyping of *tvpfp-1* mutants. Sequence in red indicate adaptors for Greengate cloning |
| LBb1 | GCGTGGACCGCTTGCTGCAACT | Genotyping mutant allele of *tvpfp-1* |
| pOS445 | accaGGTCTCaATTGAATTCTCCACTTACATCGAgttttagagctagaaatagcaag | Cloning of pJF1031-TVPFP to generate *tvpfp-2*. Red indicates the sequence of the sgRNA |
| pOS446 | tggtGGTCTCtAAACTTAAAGCTGCTTAAGAATGcaatctcttagtcgactctacc | Cloning of pJF1031-TVPFP to generate *tvpfp-2*. Red indicates the sequence of the sgRNA |
| pOS458 | CGTGATCCTAAGAACAGATAGACACG | Genotyping *tvpfp*-2 mutants |
| pOS459 | GGAGAGCTGGAATGGCTAGTATC | Genotyping *tvpfp*-2 mutants |
| pOS538 | GGCAATTACCTGTACGGGTTGACG | Primer forwards for RT-PCR. Amplifies the last exon of TVPFP |

| | | |
|---|---|---|
| pOS539 | AATGCAGTAACCAACAGACCCACC | Primer reverse for RT-PCR. Amplifies the last exon of TVPFP |
| EF1a_For | TCACCCTTGGTGTCAAGCAGA | Primer forward for RT-PCR. Amplifies the reference gene encoded by the locus AT1G07930 (ELONGATION FACTOR 1 ALPHA) |
| EF1a_Rev | CAGGGTTGTATCCGACCTTCTT | Primer reverse for RT-PCR. Amplifies the reference gene encoded by the locus AT1G07930 (ELONGATION FACTOR 1 ALPHA) |

**Table S2.** Antibodies used in this study.

| Antibody | Company/ Catalog number (Cat.no) | Dilution |
|---|---|---|
| anti-PsaA | Agrisera /Cat.no. AS06 172 | 1: 1,000 |
| anti-PsaD | Agrisera /Cat.no. AS09 461 | 1: 2,500 |
| anti-PsbD | Agrisera /Cat.no. AS06 146 | 1: 5,000 |
| anti-PsbC | Agrisera /Cat.no. AS11 1787 | 1: 1,000 |
| anti-AtpB | Agrisera /Cat.no. AS05 085 | 1: 10,000 |
| anti-ATPC | Agrisera /Cat.no. AS08 312 | 1: 1,000 |
| anti-PetA | Agrisera /Cat.no. AS20 4377 | 1: 2,000 |
| anti-LHCA1 | Agrisera /Cat.no. AS01 005 | 1: 5,000 |
| anti-LCHB2 | Agrisera /Cat.no. AS01 003 | 1: 5,000 |
| anti-RbcL | Agrisera /Cat.no. AS03 037 | 1: 5,000 |
| anti-GFP | Sigma-Aldrich/ Cat. no. G1546 | 1: 1,000 |
| anti-rabbit HRP | Agrisera /Cat.no. AS09 602 | 1: 5,000 |
| anti-mouse HRP | Agrisera /Cat.no. AS10 964 | 1: 5,000 |

### Supplementary references

1. J. Waese *et al.*, ePlant: Visualizing and Exploring Multiple Levels of Data for Hypothesis Generation in Plant Biology. *Plant Cell* **29**, 1806–1821 (2017).
